# Robust Methods For Quantifying Neuronal Morphology And Molecular Signaling Reveal That Psychedelics Do Not Induce Neuroplasticity

**DOI:** 10.1101/2024.03.04.583022

**Authors:** Umed Boltaev, Hyun W. Park, Keaon R. Brown, Maya Delgado, Jorryn Wu, Brianna N. Diaz-Pacheco, Maria Botero Pinzon, Keer He, Erin Ahern, Nina Goldshmid, Eleanor H. Simpson, Dalibor Sames

## Abstract

Induction of neuroplasticity has become the dominant explanatory framework for the rapid and sustained therapeutic effects of classic psychedelics. Within this broad concept, examination of morphological neuronal plasticity, such as dendritic arbor growth, is widely used to assess the neuroplasticity effects of classic and novel psychedelics. At the molecular level, it has been reported that serotonergic psychedelic compounds mediate dendritogenesis via the master molecular regulator of plasticity, TrkB, either directly via BDNF/TrkB signaling potentiation or indirectly through 5-HT2A receptor. To examine these hypotheses in detail, we developed a robust multimodal screening platform for unbiased, semi-automated quantification of cellular morphology and multiplex molecular signaling in the same cortical neurons. We found that in widely used primary neuronal cultures psychedelics do not directly modulate TrkB receptor or BDNF-TrkB signaling. We also found 5HT2a receptor gene expression and functional receptor levels are low, and psychedelics do not induce morphological growth, in contrast to significant dendritogenesis elicited by BDNF. Our results challenge recently published results in the field and indicate a need for rigorous experimental methods to study morphological manifestations of neuroplasticity effects induced by clinically used and experimental therapeutics.

## Introduction

Neuroplasticity is a broad term capturing a wide range of processes in the nervous system that take place in response to the changing environment. Induction of certain forms of beneficial neuroplasticity has become a global neurobiological explanatory model of our times for the effects of neuro-therapeutics such as SSRI antidepressants and psychedelics^1,2^. Similarly, molecular targets associated with neuroplasticity have garnered attention as potential avenues for therapeutic intervention^3^. In this broad context, Brain-Derived Neurotrophic Factor (BDNF) and its receptor, TrkB, have emerged as one of the central signaling hubs^4,5^. However, the design of brain penetrant, small molecule therapeutics that directly target BDNF-TrkB signaling pathways has not been achieved, underscoring the need for alternative strategies such as indirect and circuit-specific modulation of TrkB receptor^6^. Currently, an alternate class of drugs are being promoted as potential plasticity-dependent therapeutics. For this purpose, psychedelic compounds have re-emerged as a promising frontier in clinical neuroscience^7–12^. The rapid and sustained therapeutic effects of psychedelics^13^, for example in mood and substance use disorders have spurred intense interest in this class of molecules and sparked an explosion of both clinical trials and non-clinical research aimed at understanding the underlying neural mechanisms. It has been suggested that psychedelics induce neuroplasticity such as dendritogenesis and synaptic remodeling via the activation of 5HT2a receptor, molecular signaling of which may crosstalk with BDNF/TrkB signaling, or independently activate the same or similar molecular response^14–17^.

In specific terms, it is well documented that BDNF-TrkB signaling initiates three major parallel signaling cascades defined by PLCγ, Akt, and Erk activation, which mediate TrkB-regulated gene expression and ultimately changes in neuronal morphology associated with the BDNF activity^18^ (Fig. 1A). It is hypothesized that the reported dendritogenesis or synaptic plasticity induced by psychedelic compounds is mediated by the overlap in molecular signaling of TrkB and 5HT2a receptors. Indeed, 5HT2a G protein signaling is mainly transduced by Gq-PLCβ^19,20^ signaling, which has been reported to also induce Erk activation^21^, and as such covers 2/3 of TrkB major signaling cascades (Fig. 1A-“1?”). An alternative hypothesis suggests that 5HT2a agonism increases BDNF expression and release which leads to BDNF-TrkB signaling (Fig. 1A-“2?”)^22,23^. Given previous reports of TrkB transactivation by other G protein-coupled receptors (GPCRs), (namely A2A^24^, D1^25^, and Oxytocin receptor^26^), we hypothesize that 5HT2a engages similar signaling cascade leading to BDNF-independent TrkB activation (Fig. 1A-“3?”)^21^. And lastly, a recent report has claimed direct interaction between psychedelic compounds and TrkB, acting as positive allosteric modulation (PAM) of the receptor (Fig. 1A-“4?”)^15^.

**Figure 1.**
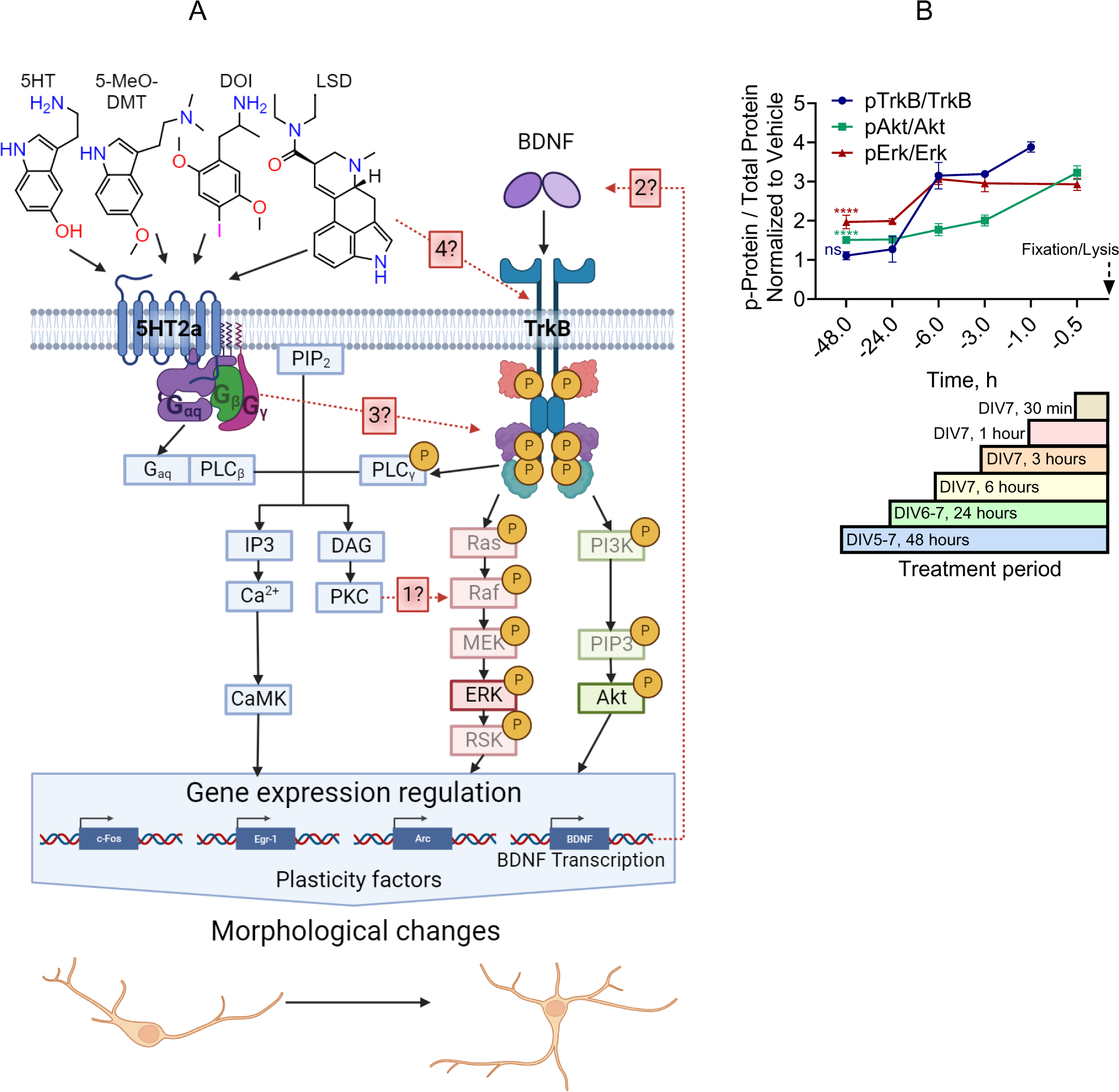
Four main molecular hypotheses for the relationship of 5-HT2a receptor, TrkB receptor, and neuronal morphological plasticity. **A.** Molecular signaling of 5HT2a and TrkB receptors. The agonism of 5HT2a receptor leads to Gq mediated activation of PLCβ, which initiates 2 parallel signaling cascades via hydrolysis of PIP2 into IP3 and DAG molecules. IP3 induces Ca^2+^ release and CaMK activation, while DAG activates PKC and, subsequently, Erk kinases, both cascades leading to gene expression regulation. TrkB activation initiates 3 major parallel signaling cascades defined by PLCγ, Erk, and Akt kinases activity and gene regulation, and subsequent morphological changes. It is hypothesized that 5HT2a activity produces morphological changes similar to TrkB activity either through the overlapping signaling cascades (IP3 and Erk), or TrkB transactivation through the unknown pathway, or BDNF expression and release. An alternative hypothesis of morphological changes induced by psychedelics proposes a direct interaction and modulation of TrkB receptor. **B.** BDNF induced TrkB, Erk, and Akt phosphorylation in rat embryonic neuronal cortical cultures (RtEN) from DIV5 to DIV7. TrkB signaling was measurable at Akt and Erk signaling molecules for at least 48h post BDNF treatment at 50 ng/ml. Data represent mean ± 95% CI from different experimental plates, two-way ANOVA, Dunnet’s multiple comparison to vehicle response, ****p < 0.0001, n=4.

To investigate these hypotheses, we developed an optical screening platform to enable a quantitative study of both morphological and molecular signaling effects in the same neuronal systems. Neuronal dendritogenesis has become a go-to method for rapid assessment of the effect of the psychedelic compounds thanks to its simplicity and scalability.^15,27–29^ Therefore, as the first step, we focused on the development of a fluorescence based dendritogenesis assay with a comprehensive experimental design to allow for quantification of morphological changes in different neuronal population and to account for different sources of biases and variations. Here we report a robust, unbiased, partially automated morphological analysis pipeline, MORPHAN (**M**orphan **O**f **R**ectified **P**aired **H**ierarchical **A**utomated **N**eurotracings) and acquisition of robust datasets and results that challenge recent claims about the molecular mechanisms of neuroplasticity induced by serotonergic psychedelics.

## Results

### Experimental Design of MORPHAN

Currently, dendritogenesis assays to study psychedelic compounds are typically performed by immunofluorescent staining of the microtubule-associated protein, MAP2, in cultured neurons at early days in vitro (DIV)^28,29^. To facilitate reconstruction of the neurons, cultures are seeded at a low density, and neurons without any overlap with other cells are manually selected for the analysis (Fig. S1). Thus, the selection of neurons is at the discretion of the experimenter. To eliminate any selection bias, we stained neurons with TdTomato fluorescent protein (TdT) through AAV infection at low titer while keeping the cultures at the same density used in molecular signaling assays^6^ allowing for sparse labeling and well separated labeled neurons. In addition to addressing the selection bias, this approach permits time series studies in which neurons are imaged before and after treatment. Such repeated, within-sample measurements provide robust datasets through internal normalization minimizing interexperimental variability (Fig. 2A). Another limitation of typical dendritogenesis assay stems from the manual nature of neuronal tracing which limits the throughput of the assay. We addressed these critical issues by developing an automated neuronal tracing algorithm for robust, unbiased, and rapid image analysis allowing for an increase in the number of neurons analyzed.

**Figure S1.**
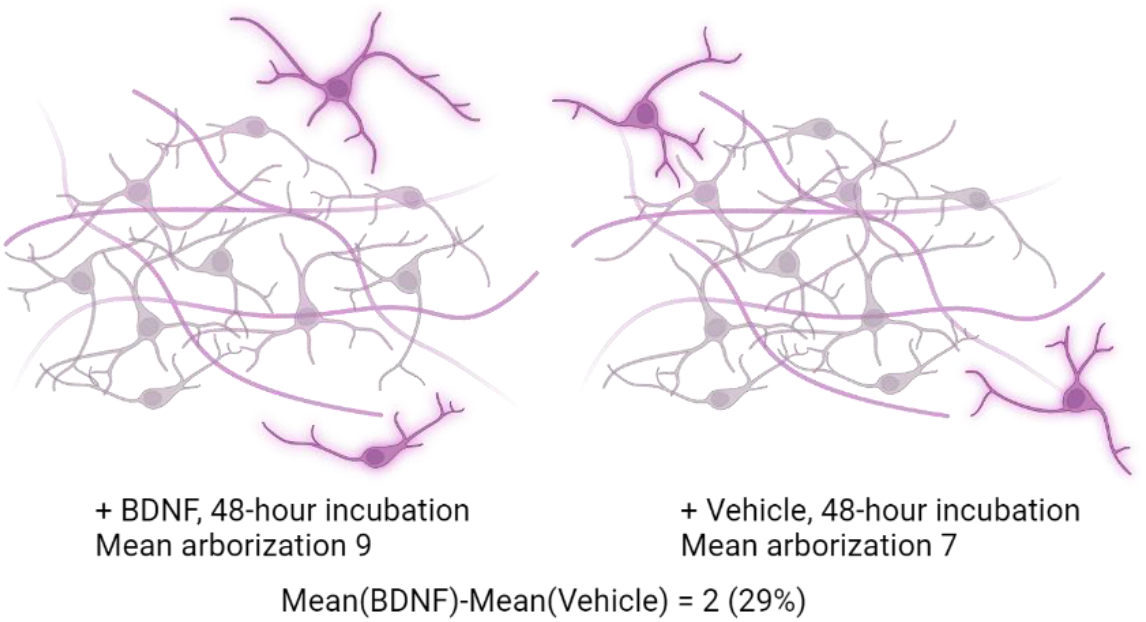
Widely used experimental design for dendritogenesis measurement. Neuronal cultures seeded at low density to achieve little overlap between the projections. After the treatment, neuron immunostained against MAP2, the isolated neurons manually selected and traced. Average arborizations are compared between the treatment groups to infer the effect of the compounds on neuronal morphology.

**Figure 2.**
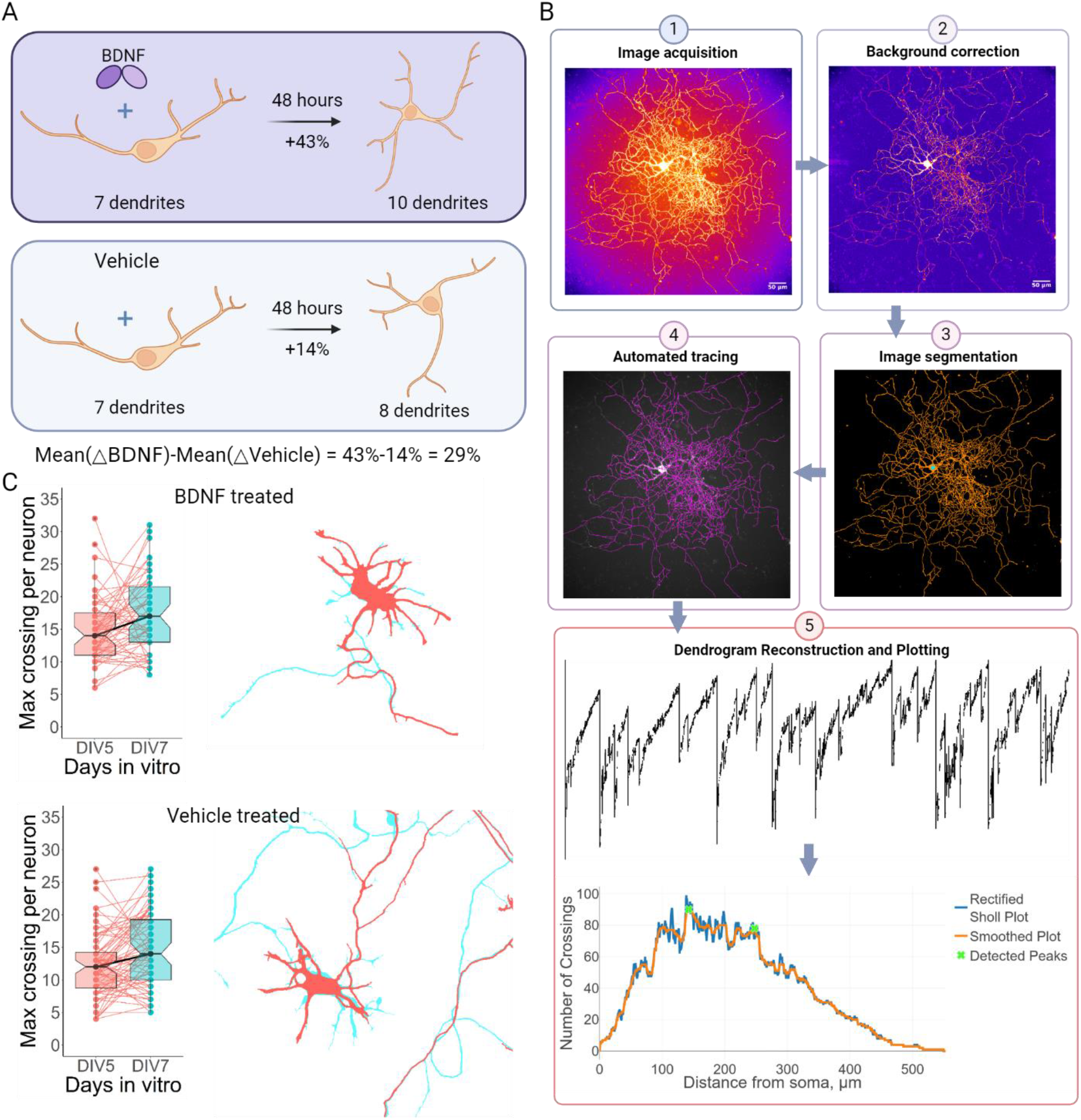
Development of a new, unbiased experimental platform for repeated imaging of neuronal cultures within treatment groups. **A.** Experimental design for dendritogenesis assay. The same cortical neurons randomly stained with TdT via AAV infection were imaged before and 48 hours after the treatment. Changes in the number of dendrites between the measurements were quantified and compared to the vehicle treated group to infer if the treatment induced a significant increase or decrease in arborization growth. Synthetic values provided for demonstration. **B.** Image analysis and data extraction pipeline of neuronal morphology. *B1*. Image acquisition of fluorescently labeled neurons. Image quality was a critical aspect in performing robust image analysis. *B2*. Background correction to account for spherical aberration and uneven illumination. A new algorithm was developed to create an accurate representation of the background profile of the given image with subsequent correction of the background. *B3*. Image segmentation. To accurately separate neurons from the background, a Weka machine learning model was developed for each dataset. Orange – neurites, cyan – cell body. *B4*. Automated tracing. APP2 plugin of Vaa3D was used to trace each neuron separately using predetermined soma locations, purple – traces overlayed over original image. *B5*. Trace straightening and plotting. Since some projections were looping, traces were straightened to avoid double counting. The straightened representation of the traces was used to create a rectified Sholl plot. Using median smoothing filter, max number of projections within 150 μm from soma were determined and used for statistical analysis. Image of TdT stained neuron on DIV12 presented for demonstration of the pipeline. **C.** Repeated measurements of neuronal arborization of rat embryonic neuronal (RtEN) cortical cultures treated with BDNF at 50ng/ml (top) or vehicle (bottom) and a representative neuron (right outlines, red – DIV5, cyan – DIV7), highlighted in black on the graphs. BDNF and vehicle treated samples increased in the maximal number of crossings by 15% (p = 0.0084, N = 55) and 20% (p < 0.0001, N = 72), respectively. The difference in the changes between the treatment groups was not significant (p = 0.88, see Fig. S1).

### Rat embryonic neuronal culture as a primary model for dendritogenesis assay

Rat embryonic neuronal (RtEN) cultures are a well optimized and a widely used in-vitro model to study both molecular signaling and morphological changes of neurons natively expressing proteins of interest. Considering BDNF as a positive control, a typical time frame for published dendritogenesis assays is 3-10 days in vitro (DIV3-10^30–32^). BDNF does not appear to affect neuronal arborization in mature cultures (beyond DIV12)^33^. To determine the optimal DIV for pretreatment imaging, we infected RtEN cultures on DIV1 with TdT under CAG promoter and observed sufficient expression for accurate tracing in a substantial number of neurons starting from DIV5.

The typical treatment period for dendritogenesis assay has been 2-3 days. To be certain that TrkB receptor is functionally expressed within this period, we determined how long TrkB receptor signaling remains active by measuring phosphorylation of TrkB, Akt, and Erk signaling molecules through ELISA and ELFI in samples treated 0.5 to 48 hours before fixing or lysing the cells on DIV7 (Fig. 1B). The activity level declined both at the receptor level and downstream targets. We found that TrkB phosphorylation was undetectable at 48 hours post treatment, while Akt and Erk were still significantly elevated in BDNF treated samples compared to vehicle controls, indicating continuous TrkB downstream signaling for 48 hours post BDNF addition. Based on these results, we chose to use DIV5 and DIV7 as the window to measure morphological changes.

### Image Analysis pipeline

MORPHAN consists of 5 major steps (Fig 2B). Image quality plays an important role in the efficiency of the analysis, so we sought to image brightly stained neurons. After the image acquisition, background correction was performed to mitigate spherical aberration, an uneven illumination introduced by the microscope hardware (Fig 2B1). For this purpose, we created a new algorithm based on a nonlinear regression model to simulate the image background, which are then subsequently used to flatten the original image and equalize the fluorescent signal (Fig 2B2). To segment the images, we used the intensity-based machine learning WEKA segmentation ImageJ plugin^34^, as it provided the most flexibility and accuracy to isolate neurons from the background (Fig 2B3). To reduce experimenter bias, we prepared training set of images with arbitrary file names so that the operators of WEKA were blinded, and a single WEKA model was generated for each dataset. The resultant binary images contained disconnection at the position with low signal, which were manually corrected by the operator blinded to the identity of the sample.

Generated binary images were subsequently subjected to automated tracing using the APP2 plugin from the Vaa3D software suite^35,36^. While the plugin accurately traced images with a single neuron as confirmed by manual tracing (Fig. 2B4, S2A-C), it did not separate the boundaries of 2 or more adjacent cells (Fig. 3D). To resolve these traces, we used the G-Cut algorithm developed specifically for this task^37^. Using G-Cut parameters for the cortical neurons, we successfully separated traces letting the algorithm assign the projection to the corresponding soma (Fig. 3D). We reasoned that human operators would not be able to correctly determine from which cell any given projections originate, so we allowed the parameters in the algorithm to make those decisions to result in consistent bias. Since the tracings generated by APP2 were dependent on the soma position, we adopted a soma-centric approach, where any given neuron was subjected to both APP2 tracing and G-Cut re-tracing (Fig. 3D). This resulted in one projection being associated with 1 or more neurons. We accepted such an outcome since we were interested in comparative studies. The pipeline can reliably trace complex projections of TdT stained neurons (Fig. 2B).

**Figure 3.**
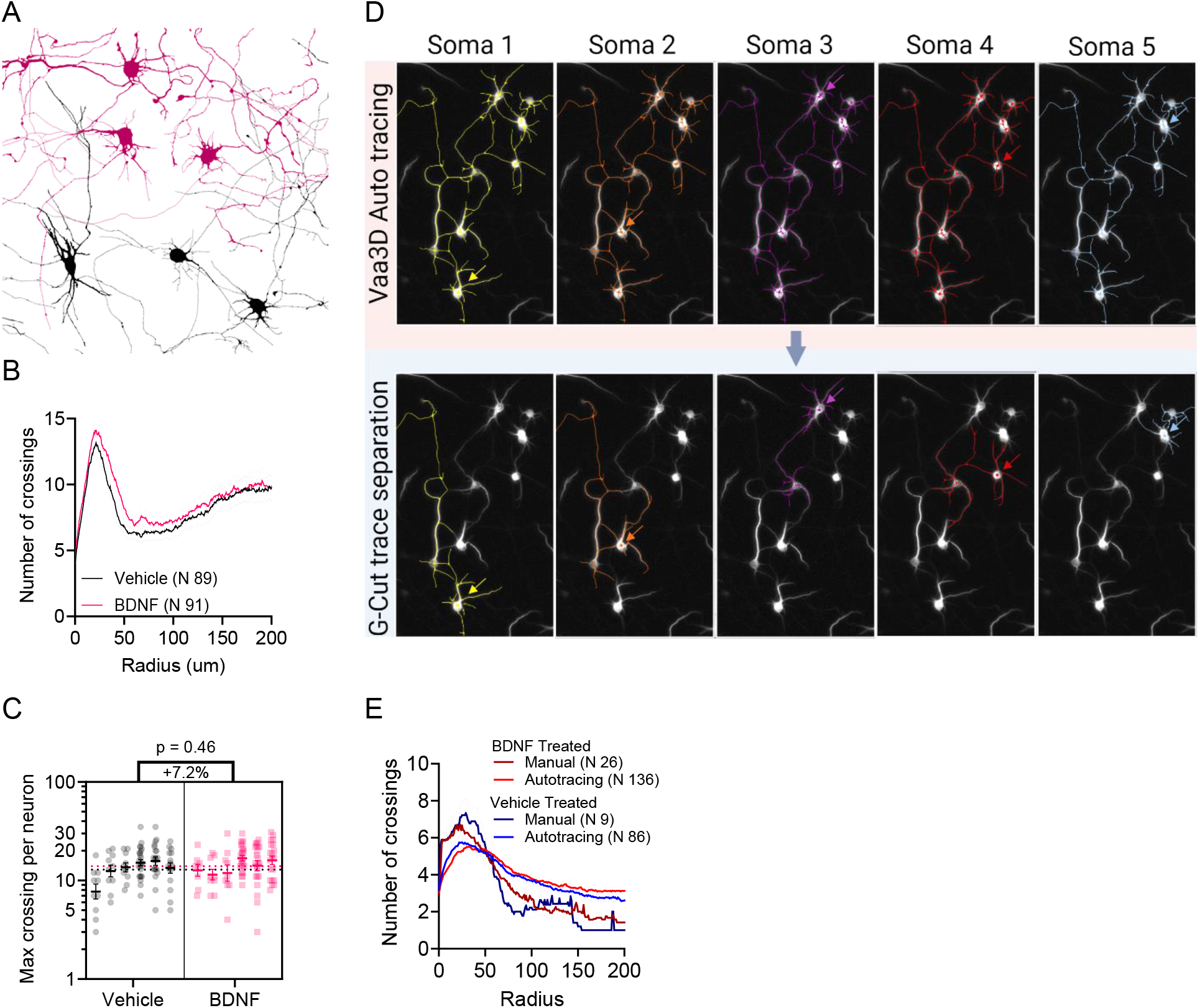
Snapshot cross-sample comparison of AAV and MAP2 stained neurons. **A.** Outlines of representative neurons on DIV7 in RtEN cortical cultures treated with BDNF at 50 ng/ml (magenta) or vehicle (black) on DIV7. **B.** Rectified Sholl plots of the neurons in A. **C.** Max number of crossings segregated by individual plates as columns. BDNF did not change the arborization of the treated neurons versus vehicle. Data represent mean ± 95% CI of back-transformed estimates derived from the generalized linear model with negative binomial distribution (deviation between the plates was insignificant, plate number = 6, p-value = 0.086). Number of traced neurons shown in B, effect size and p-value in C. **D.** Trace separation of MAP2 stained neurons. To quantify as many neurons as possible and separate the traces, G-Cut algorithm was applied which assigned the projection to the corresponding soma based on parameters for cortical neurons. Each neuron was run through Vaa3D tracing and G-Cut separation algorithms separately. Arrows point to the specific soma. **E.** Sholl plots of BDNF and vehicle treated RtEN cortical cultures derived from tracing of MAP2 stained neurons. Both manual and autotracing method found no significant difference between the treatment groups.

To address the shortcomings of classical Sholl analysis, such as double counting of the looping projections, and to obtain realistic representations of neuronal arborization from the point of view of the soma, we used dendrograms as the representation of neuronal arborization^38^. The rectified Sholl plot was then generated by counting the number of projections at a given distance starting from soma (Fig. 2B5). This algorithm provided smoother plots (Fig. S2D). To compare the accuracy of auto tracing with experimenter tracing, we manually traced a subset of the acquired images and compared them to the outcome of the pipeline. Generated rectified Sholl plots were similar between manual and automated traces (Fig. S1C-E).

**Figure S2.**
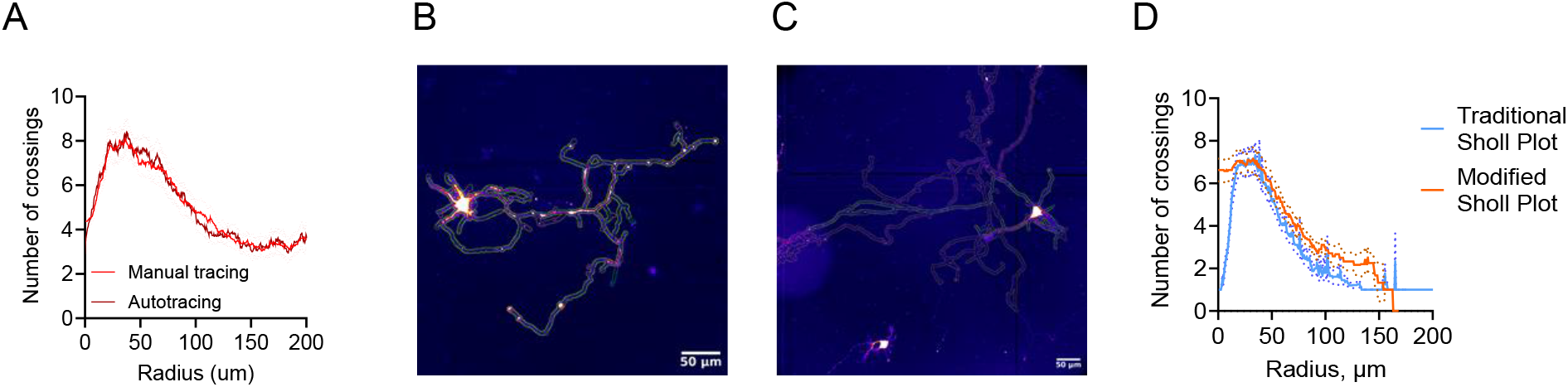
Pipeline evaluation. **A**. Comparison of average straightened Sholl plots of manually traced and auto traced neurons stained with TdT. Both tracing methods produced similar results. Mean (solid line) ± SEM (dashed line). **B**. A comparison of traces with the least difference between manual and autotracing. “Fire” pseudo color – image with TdT stained neuron, green overlay – manual tracing, orange overlay – auto tracing. **C**. A comparison of traces with the most difference between manual and auto-tracing. Color assignment is the same as in B. **D**. Comparison of traditional Sholl plot and straightened Sholl plot. Traditional plots contained a number of local maxima in the form of “spikes”, while straightened plots were smoother. Mean (solid line) ± SEM (dashed line).

While there are many morphological features to be used for statistical analysis^39–41^, we decided to focus on a single measurement of dendritic arborization which we defined as the maximal number of crossings (NMax) within a 100 μm radius from the soma. To measure the NMax, we smoothed the Sholl plot of the individual neuron to remove the “spikes” and found the peak within 100 μm (Fig. 2B5). Given that NMax is a count value and to factor in variations between the seedings, we identified appropriate statistical models for the specific comparisons (see Methods)^42^.

### Using MORPHAN to Quantify the Effects of BDNF And Psychedelic Drugs on Dendritogenesis

Using the MORPHAN pipeline, we examined RtEN cortical cultures for morphological response to BDNF treatment. The cultures were imaged prior to treatment on DIV5 and randomly assigned to treatment groups. The assignment and treatment were performed by an experimenter not involved in imaging and image processing to eliminate selection bias. The same neurons were re-imaged and analyzed after 48 hours. For within-sample analysis, neurons were manually assigned an ID and matched across imaging sessions. Between DIV5 and DIV7, arborization increased by 20% and 15% in vehicle and BDNF treated samples respectively (Fig. 2C). The difference in growth between the samples was not significant (−4.3%, p = 0.5, Fig. S3A). We were surprised to find no effect of BDNF on growth, so we performed a population comparison of neurons on DIV7 (a single timepoint between-sample comparison). We found a significant increase in neuronal arborization induced by BDNF treatment (+17%, p = 0.014, Fig. S3B). When we compared neuronal arborization at DIV5, however, we observed significant differences between the groups at the baseline before treatment (+23%, p = 0.0014) which could explain the observed post-treatment difference. Moreover, when we included all the TdT stained neurons imaged on DIV7, we found no significant effect of BDNF (+7.2%, p=0.27, Fig 3A-C). Given these observations, we concluded the significant effect we observed with the pre-imaged population of TdT stained neurons on DIV7 was a false positive, illustrating the importance of repeated within-sample comparisons.

**Figure S3.**
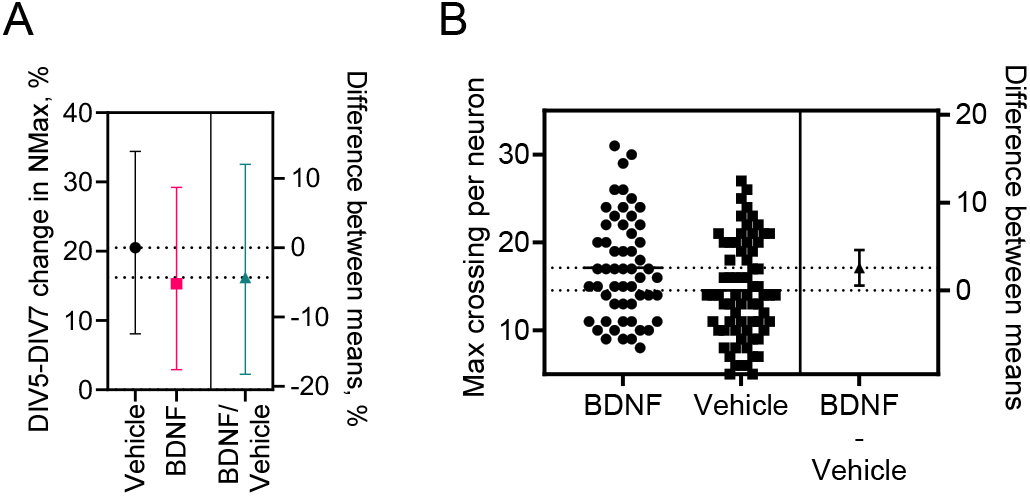
BDNF dendritogenesis effect in RtEN. **A**. Changes in neuronal arborization between DIV5 and DIV7 vehicle or BDNF treated samples, and the difference between the groups (Fig. 2C). Data represent mean ± 95% CI of back-transformed estimates derived from generalized linear mixed model with Poisson distribution, N in vehicle group = 72, N in BDNF group = 55. **B.** Max number of crossings of neurons in Fig. 2C, DIV7. BDNF was found to significantly increase neuronal arborization (p = 0.014). Data represent mean ± 95% CI of back-transformed estimates derived from the generalized linear model with negative binomial distribution (deviation between the plates was insignificant, plate number = 6, p-value = 1). N in vehicle group = 72, N in BDNF group = 55.

Given previous reports ^30–32^, we were surprised to find that BDNF did not enhance growth. Therefore, to verify our findings, we used the typical immuno-stain, MAP2, to test the effect of BDNF treatment. We manually traced a subset of neurons and used MORPHAN pipeline on the whole dataset (Fig. 3D-E). We again found that BDNF did not significantly change neuronal arborization measured through either manual or automated tracing. RtEN cultures were not affected by BDNF treatment most likely due to the robust growth supported by the neurobasal medium alone, which masked the dendritogenic effect of TrkB activation.^43^ Therefore, we concluded that without a positive control these neuronal cultures cannot reliably be used to study the effect of psychedelics or other bioactive compounds on dendritogenesis and neuronal morphology.

### Psychedelics do not activate or potentiate TrkB receptor

RtEN cortical cultures are, however, well suited to study TrkB molecular signaling (Fig. 1B).^6^ To this end we tested a representative set of psychedelics for TrkB, Akt, and Erk activation as agonists in time course experiments. While BDNF induced TrkB, Akt, and Erk activation in a time and dose-dependent manner, serotonin, DOI, 5-MeO-DMT, and LSD did not produce any activation of these signaling molecules at any time point (Fig. 4A). These results demonstrate that psychedelic compounds do not activate TrkB as agonists, in the absence of BDNF, or induce downstream pathways Akt and Erk in RtEN.

**Figure 4.**
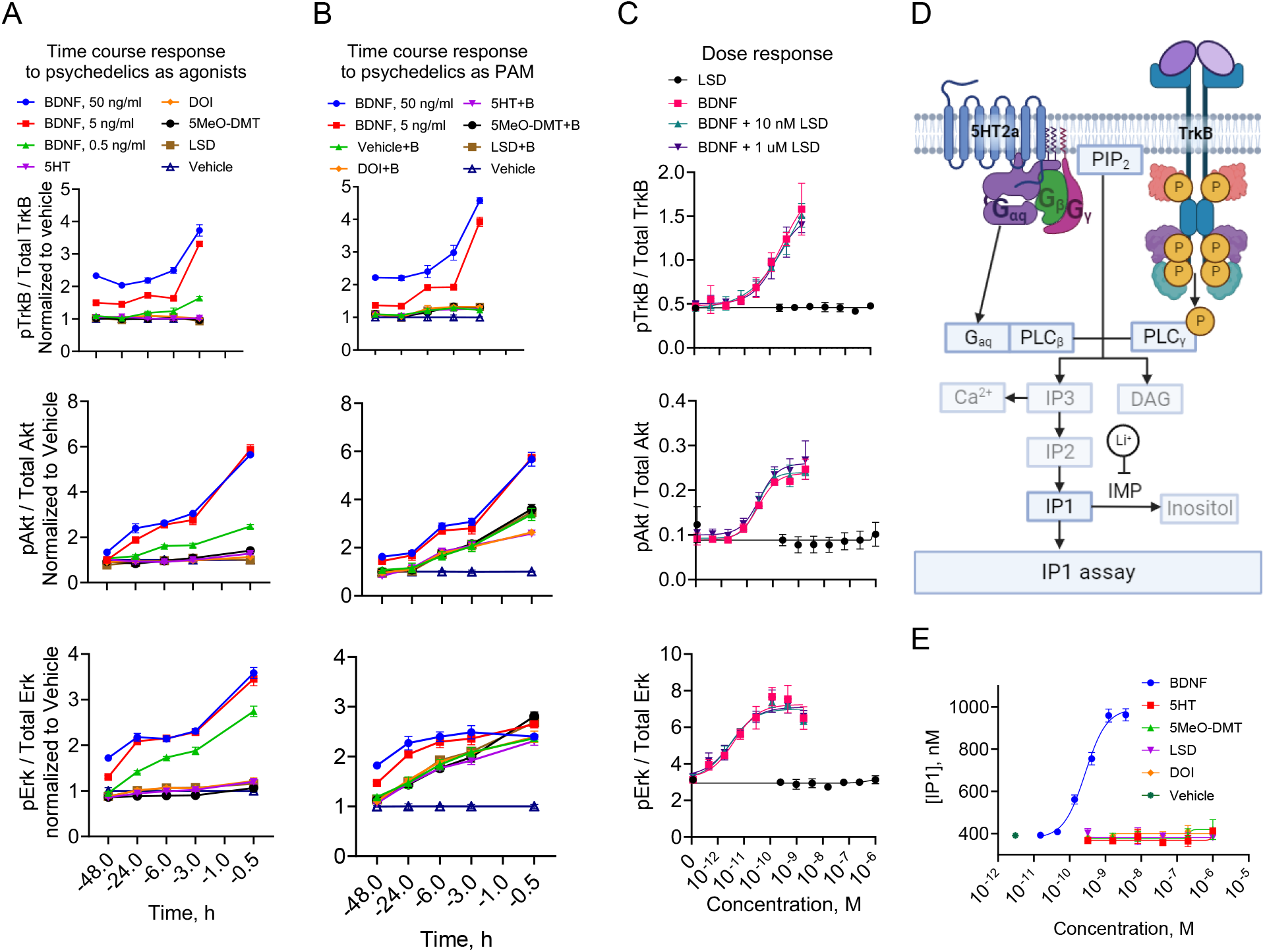
Psychedelics do not modulate TrkB receptor in RtEN cultures. **A.** TrkB, Akt, and Erk activation induced by compounds in RtEN cortical cultures. Cultures were treated on DIV5 for 48h, DIV6 for 24h, or DIV7 for 6h, 3h, and 30min (Akt, Erk) or 1h (TrkB) before fixation. The concentration of DOI, 5HT, 5MeO-DMT, and LSD were at 1 μM, BDNF concentration at 50 ng/ml, 5 ng/ml, 0.5 ng/ml. Compounds did not activate TrkB or its downstream targets at any time point of the incubation. **B.** Psychedelics as positive allosteric modulators of TrkB. RtEN cortical cultures were treated as in A with compounds co-incubated with BDNF at 1 ng/ml (indicated as “+ B” in the legend, EC6 at TrkB, EC50 at Akt, EC85 at Erk). Psychedelics did not demonstrate PAM activity at any time point tested. **C.** Dose response of LSD, BDNF, and BDNF in the presence of LSD at 10 nM and 1 μM as PAM at TrkB, Akt, and ERK in RtEN cortical cultures. Neurons were treated for 3h on DIV7. LSD did not induce activation of TrkB signaling in agonistic, nor potentiated BDNF activity. Data represents mean ± SEM. **D.** IP1 assay principle. Both 5HT2a and TrkB induce IP3 signaling through PLCβ and PLCγ, respectively. Treatment of cell cultures with LiCl leads to inhibition of IP1 hydrolysis. Cells were lysed for subsequent IP1 assay. IMP = Inositol monophosphatase. **E.** IP1 accumulation in RtEN cortical cultures induced by BDNF and psychedelics for 1h on DIV5. BDNF induced IP3 signaling (EC50 = 165 pM, or 5 ng/ml), psychedelics did not produce any IP1 accumulation. Data represents mean ± SEM of interpolated values.

To examine the recently reported TrkB potentiation by psychedelics,^15^ we tested the compounds in the presence of BDNF at 1ng/ml, which corresponds to EC6 at TrkB, EC50 at Akt, and EC80 at Erk. Neither serotonin, DOI, 5-MeO-DMT, nor LSD produced any potentiation of these signaling molecules at any time point (Fig. 4B). To probe further, we examined the effect of 2 concentrations of LSD at which it has been reported to bind to the receptor^15^ (10 nM and 1 μM) on the dose response of BDNF at TrkB, AKT, and Erk, to find no effect of LSD (no shift in EC50 values of BDNF, Fig. 4C). We also tested the effect of LSD on NT3 dose response, a TrkB partial agonist, to find no changes in the EC50 values of this neurotrophic factor (Fig.S4A). Lastly, we also tested the effect of LSD on BDNF dose response in CHO-TrkB cells, once again, to find no effect (Fig.S4B). These results indicate that psychedelics do not functionally interact with TrkB or the downstream signaling kinases.

**Figure S4.**
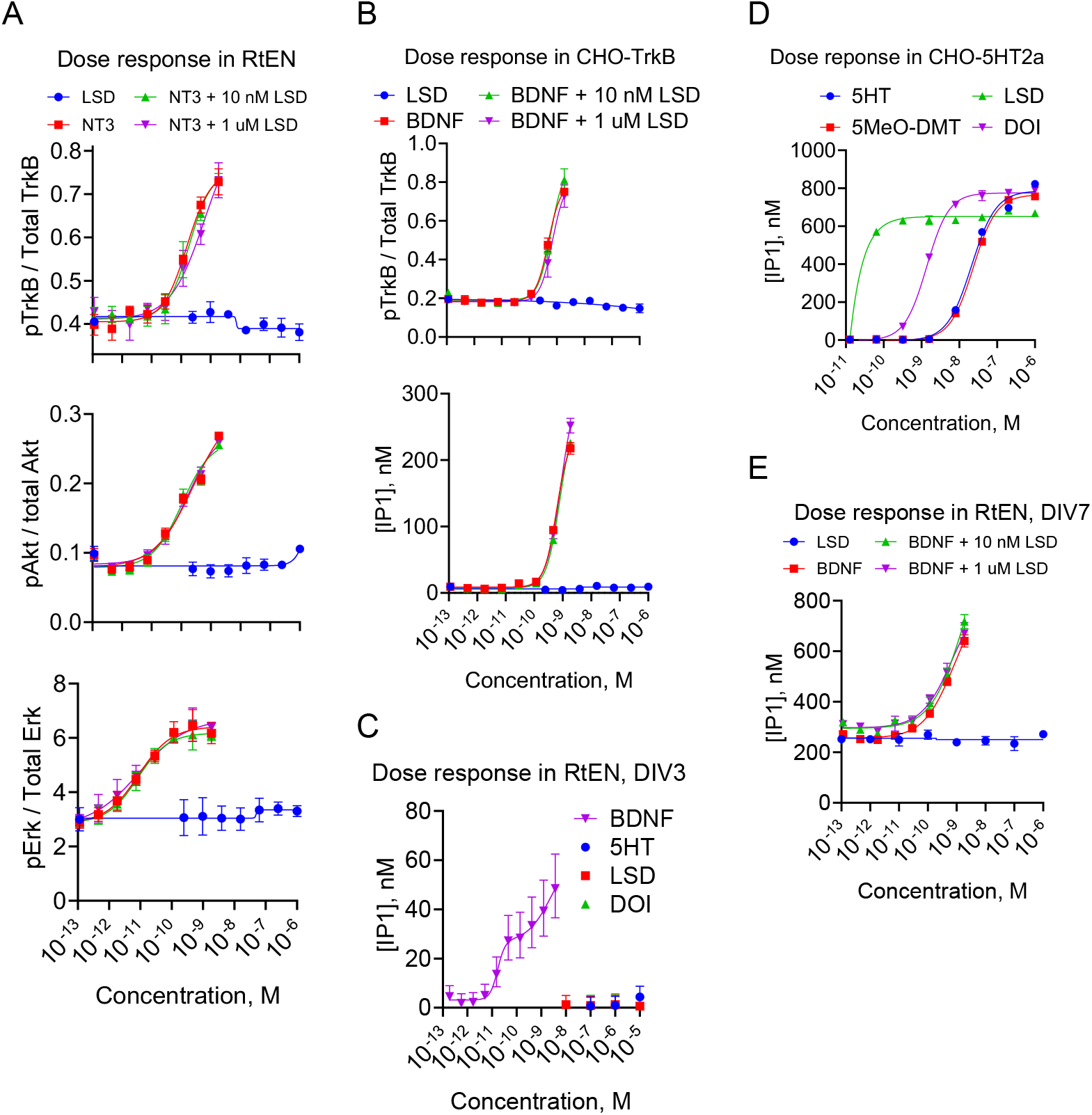
Psychedelics do not modulate TrkB in cortical neurons or TrkB-CHO cells, nor activate IP3 pathway in cortical neurons. **A**. Dose response of LSD, NT3, and NT3 in the presence of LSD at 10 nM and 1 μM as PAM at TrkB, Akt, and ERK in RtEN cortical cultures. Neurons were treated for 3h on DIV7. LSD did not potentiate TrkB partial agonist, NT3, signaling. **B**. Dose response of LSD, BDNF, and BDNF in the presence of LSD at 10 nM and 1 μM as PAM at TrkB and IP3 in CHO-TrkB cell cultures. LSD did not potentiate BDNF in the overexpressed system. Data represents mean ± SEM. **C**. IP1 accumulation in RtEN cortical cultures induced by BDNF and psychedelics for 1h on DIV3. BDNF induced IP3 signaling in a biphasic dose dependent manner, psychedelics did not produce any IP1. Data mean±95%CI of interpolated values for demonstration purpose. Nonlinear fit to determine EC50 values were performed on HTRF values. **D**. IP1 accumulation in CHO-5HT2a cell line induced by psychedelics for 1h. Psychedelics were active at 5HT2a in the overexpressed system. 5HT EC50 = 5 nM, 5MeO-DMT EC50 = 5 nM, DOI EC50 = 0.4 nM, LSD EC50 < 64 pM. Data represents mean ± SEM of interpolated values for demonstration purposes. Nonlinear fit to determine EC50 values were performed on HTRF values. **E**. Dose response of LSD, BDNF, and BDNF in the presence of LSD at 10 nM and 1 μM as PAM at IP3 signaling in RtEN cortical cultures. Neurons were treated for 3h on DIV7. LSD did not potentiate BDNF signaling at IP3.

### 5HT2a is not functionally expressed in rat embryonic neuronal cultures

The lack of Erk activation indicated that either 5HT2a is not expressed in RtEN cultures or it did not activate Erk signaling pathway^21^. Since it has been reported that 5HT2a is expressed in the RtEN cultures^14,17,44–47^, we sought to develop an assay that could test its functional expression. The prominent BRET assays ^20,48–50^ developed over the last decades rely on the co-transfection of 3 or 4 plasmids into competent cell lines. In our experience, neurons do not demonstrate such competence, necessitating the use of AAV infection instead of lipid mediated transfection. As such we researched assays that could measure downstream signaling molecules. Given that 5HT2a activation leads to IP3 and Ca^2+^ signaling, we adapted IP1 assay to neuronal cultures^51^. The assay is based on accumulation of the IP3 derivative, IP1, through the inhibition of Inositol monophosphatase with LiCl (Fig. 4D). As such, we pre-treated the cells with LiCl for 5 min before adding the compounds. To measure the accumulation of IP1, we lysed the cells and added detection components of the IP1 assay kit. Initially, we tested psychedelics’ activity in 5HT2a-expressing CHO cells. 5HT and 5-MeO-DMT produced full agonism with the same potency of EC50 at 5 nM, while DOI appeared to be 10 times more potent full agonist (EC50 = 0.4 nM). LSD activity in our preparation produced full agonism with EC50 < 64 pM (Fig. S4D). Given that TrkB also induces IP3 signaling, we tested the assay in CHO-TrkB cells. BDNF treatment produced IP1 accumulation with the same potency as at TrkB receptor level indicating no signal amplification (Fig. S4B). LSD did not activate IP3 signaling nor TrkB potentiation in CHO-TrkB demonstrating specificity to 5HT2a receptor. These results confirm that the adapted assay produced robust signal derived from the activation of both 5HT2a and TrkB receptors. We then tested RtEN cultures at DIV3 and DIV5 for IP3 signaling induced by BDNF and psychedelic compounds. We found that BDNF induced IP1 accumulation in a dose dependent manner (EC50 173 pM or 4.6 ng/ml), while psychedelic compounds did not induce IP3 signaling at any tested concentrations (Fig. 4E, S4C). These results indicate very low/no functional levels of 5HT2a receptor present in these neurons.

### BDNF Induces Dendritogenesis in cultured mouse Neurons

Since low functional expression of 5HT2a and lack of suitable transgenic rat models prevent using RtEN cultures to analyze morphological changes resulting from 5HT2a activation by psychedelics, we investigated alternative primary cultures. Due to the availability of a wide variety of mouse transgenic models^52^ that can be used to selectively label neuronal populations expressing genes of interest, we decided to use mouse postnatal (MsPN) cortical cultures. We tested various seeding procedures and culture conditions to obtain high yields of growing neurons (see Methods). Using our MORPHAN pipeline, we examined MsPN cortical cultures for morphological response to BDNF treatment. In neurons seeded in standard medium and treated with vehicle, we found no significant changes in dendritic morphology between DIV5 to DIV7(Fig. 5A-B). In contrast, BDNF treatment produced significantly more projections showing a 60% increase over the control group demonstrating that MsPN cultures can be used to quantify dendritogenesis.

**Figure 5.**
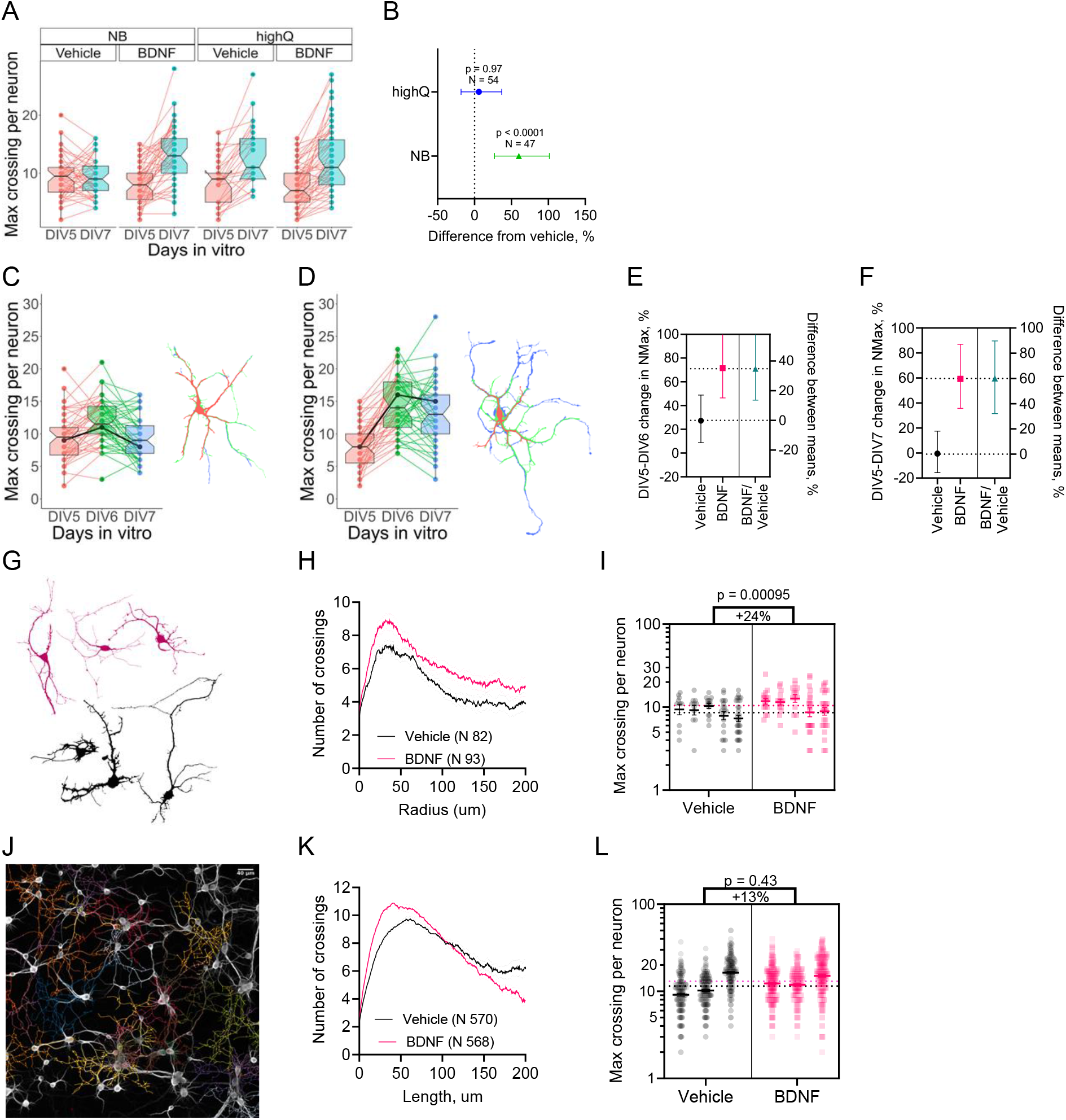
BDNF induces dendritogenesis in mouse postnatal (MsPN) cortical cultures. **A.** Arborization growth of mouse postnatal (MsPN) cortical cultures seeded in standard neurobasal medium and neurobasal medium with high glutamine treated with vehicle or BDNF (50 ng/ml). The max crossings were significantly increased in BDNF treated groups and in highQ medium. BDNF and highQ effect doesn’t appear to be additive. NB-Vehicle: - 0.45%, p = 1, N = 48; NB-BDNF: +59%, p < 0.0001, N = 47; highQ-Vehicle: +55%, p < 0.0001, N = 29; highQ-BDNF: +59%, p < 0.0001, N = 54. **B.** Changes in neuronal arborization between DIV5 and DIV7 of conditions in A. A significant increase of 60% in neuronal arborization induced by BDNF treatment was observed only in standard medium. **C.** Time course graph of arborization changes of neurons in MsPN cortical cultures in standard medium treated with a vehicle. A representative neuron shown on right (outlines, red – DIV5, green – DIV6, blue – DIV7), the corresponding data point highlighted in black on the graph. **D.** BDNF treated cultures similar to C. **E-F.** Changes of neuronal arborization measured from C and D. Vehicle treated neurons demonstrated 27% (p-value = 0.00013) increase in arborization from DIV5 to DIV6, while on DIV7 return to the arborization level that of DIV5 (−0.4%, p-value = 0.95). BDNF treatment increased the number of neurites more than vehicle by 35% (p-value = 0.0008) and by 60% (p < 0.0001) 24h and 48h post treatment, respectively. Samples before BDNF treatment were the same as vehicle treated samples (−10%, p-value = 0.17). Data in B, E, and F represent mean ± 95% CI of back-transformed estimates derived from the generalized linear mixed effect model with Poisson distribution. N in E and F: vehicle group = 48, BDNF group = 47. **G.** Outlines of the representative neurons treated with BDNF (magenta) or vehicle (black) stained with TdT, imaged on DIV7. **H-I.** Rectified Sholl plot (H) and Max crossing measurements (I) of the TdT stained neurons at DIV7. Population comparison showed 24% increase in max crossing in cultures treated with BDNF. Data represent mean ± 95% CI of back-transformed estimates derived from the generalized linear model with negative binomial distribution (deviation between the plates was insignificant, plate number = 5, p-value = 0.49). N and p values are shown on the graph. **J.** Representative image and auto-tracings of MAP2 stained neurons in MsPN cortical cultures on DIV8. **K-L.** Straightened Sholl plot (K) and Max crossing measurements (L) of neurons in J. Population comparison showed a non-significant increase of 13% (p=0.43) in max crossing in cultures treated with BDNF. Data represent mean ± 95% CI of back-transformed estimates derived from the generalized linear mixed effect model with negative binomial distribution (deviation between the plates was significant, plate number = 3, p-value < 0.0001). The number of traced neurons and p values are shown on the graph.

Interestingly, we serendipitously found that GlutaMax, a glutamine supplement in the form of glutamine-alanine dipeptide, influenced neuronal growth. Increasing the concentration of GlutaMax to 2 mM (highQ medium) promoted neuronal arborization in control samples. As a result, highQ medium masked the dendritogenic effects of BDNF (Fig. 5A-B). Neurons seeded in highQ media demonstrated similar increase of their arbor in both treatment groups (BDNF and Vehicle) demonstrating no additive effect of the two treatments. This observation indicated that the dendritic growth of neurons in primary culture was sensitive to medium composition. Increased GlutaMax treatment can therefore be used as small-molecule positive control in dendritogenesis assays.

Detailed examination of TdT labeled cultures in standard medium revealed that the arborization in vehicle treated cultures increased on DIV6 but returned to pre-treatment levels by DIV7 (Fig. 5C-D). In contrast, BDNF treated neurons demonstrated increases in arborization at DIV6 that were larger than the vehicle treated groups and were retained on DIV7. Since vehicle treated neurons reduced their arbor and BDNF treated samples sustained it, the overall effect size of BDNF on DIV7 was larger than on DIV6 (Fig. 5E-F, DIV6-DIV5 +34.8%, p=0.0008; DIV7-DIV5 +60%, p < 0.0001).

A between-sample comparison of rectified Sholl plots revealed that BDNF treated samples have larger arborization than vehicle treated samples at DIV7 (Fig 5G-H). Comparison of NMax showed a 24% increase (p = 0.00095) in neuronal arborization in BDNF treated cultures (Fig 5I). To increase the sample size and compare the outcome to our data from TdT labeling, we also examined MAP2 stained neurons for between-sample effects. To this end, the cultures described above were fixed and immunostained on DIV8. Using the same algorithm and removing traces shorter than 100 μm, we increased our sample size to more than 500 neurons per group. The Sholl plot of BDNF treated neurons showed an increase in arborization (Fig. 5J-K), however, when we factored in variation between the plates within each treatment group, the 13% increase in neuronal arborization fell short of significance (p=0.43, Fig. 3L).

A power analysis of the data showed that for either within-subject or groupwise comparisons, to achieve 80% power at effect sizes of 60%, 20% and 10% we needed to analyze 17-20, 100, and 600 neurons per treatment group respectively. We therefore concluded that the best approach to quantify morphological changes was a within-subject normalization of TdT stained neurons from DIV5 to DIV7, as it produced the largest effect size and no random effect produced by the independent seedings.

### Psychedelics do not induce dendritogenesis in mouse postnatal cultures

Following the same protocol described above, we treated MsPN cortical cultures with BDNF at 50 ng/ml, 5HT, 5MeO-DMT, DOI at 1 μM, and LSD at 500 nM. In standard growth conditions, BDNF increased neuronal arborization by 30% compared to vehicle treatment while treatment with the 5HT2a agonists produced no significant changes (Fig. 6A-B). Moreover, we observed a trend for 5MeO-DMT to reduce the growth by 16% (p = 0.09, N = 40).

**Figure 6.**
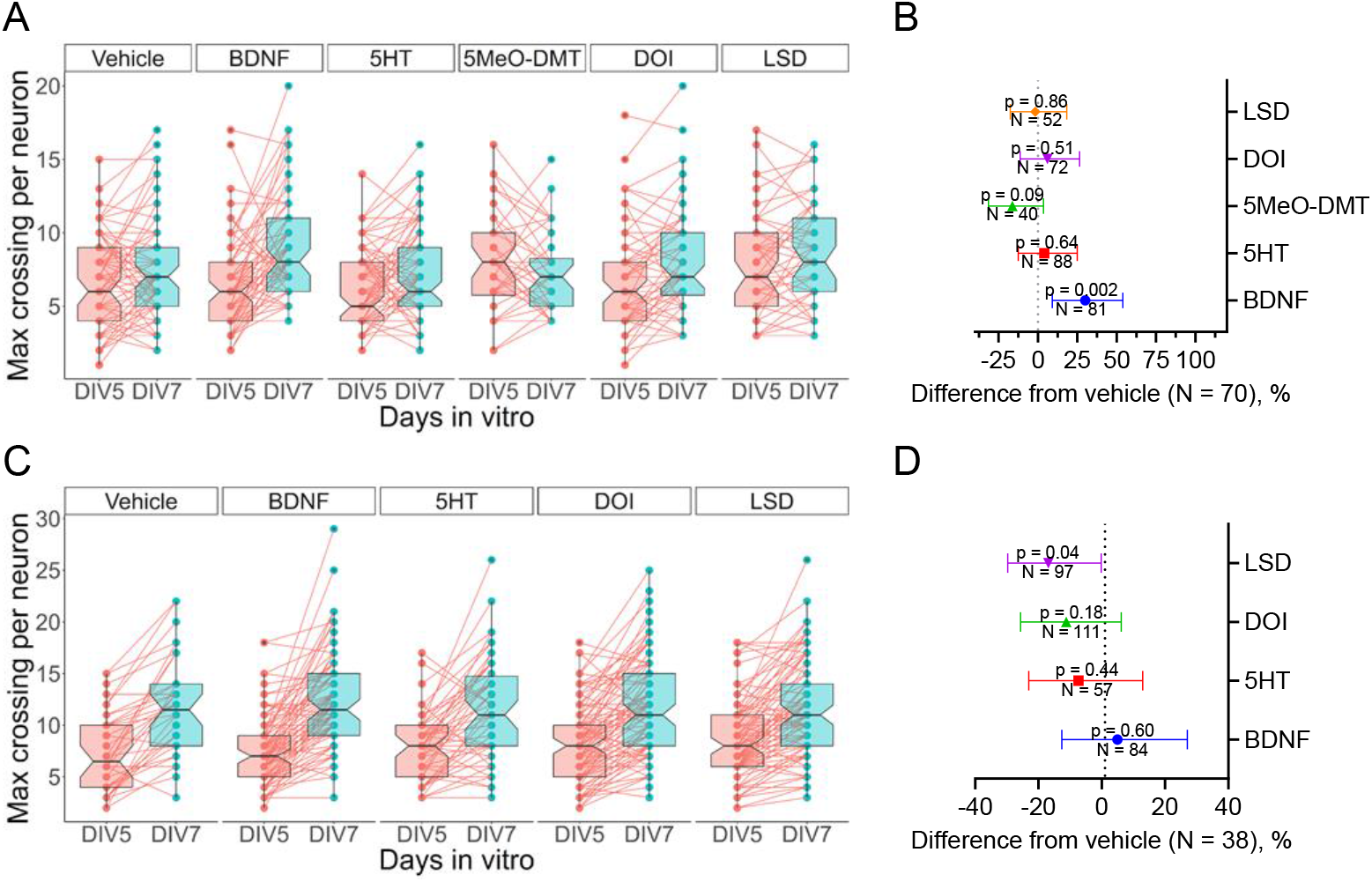
Psychedelics do not increase dendritogenesis in MsPN cortical cultures. **A.** Arborization growth of MsPN cortical cultures treated with 50ng/ml of BDNF or 1 μM of 5HT, 5MeO-DMT, DOI, or 500nM of LSD. **B.** Analysis of changes in neuronal arborization between DIV5 and DIV7 of cultures in A. A significant increase in neuronal arborization was observed only in cultures treated with BDNF, +30%. **C.** Arborization growth of MsPN cortical cultures seeded in highQ medium and treated with 50ng/ml of BDNF or 1 μM of 5HT, DOI, or LSD. **D.** Changes in neuronal arborization between DIV5 and DIV7 of cultures in C. A significant decrease in neuronal arborization growth was observed in cultures treated with LSD, −18%. Data represent mean ± 95% CI of back-transformed estimates derived from the generalized linear mixed effect model with Poisson distribution. Number of traced neurons and p values are shown on the graphs.

In highQ medium, as expected, BDNF did not increase dendritic growth more than vehicle (Fig 6C-D). Meanwhile, LSD reduced growth by 18% (p = 0.04) and we also observed a trend for reduced growth in 5HT and DOI treated samples. Overall, we found no evidence for dendritogenic effects of psychedelics in MsPN cultures in the randomly stained neurons.

Given the lack of effect of psychedelics, we tested if 5HT2a receptors are active in MsPN cultures. Using IP1 assay, we determined that BDNF, but not psychedelics, induced the accumulation of IP1 (Fig. S6D), suggesting that the receptors are either not functional or not present in the cultures. Because there are no available antibodies sensitive or selective enough to reliably report 5HT2a receptor protein level (see SM and Fig. S5), we performed single molecule fluorescent in-situ hybridization (smFISH) to quantify the number of neurons that express 5HT2a mRNA molecules (Fig. S6A-C). Specificity of the smFISH probes was confirmed in cells overexpressing mouse 5HT2a receptor (Fig. S6A). In the primary MsPN cultures, we found that 11% of all cells and only 6% of randomly TdT stained cells expressed 5HT2a mRNA molecules on DIV7.

**Figure S5.**
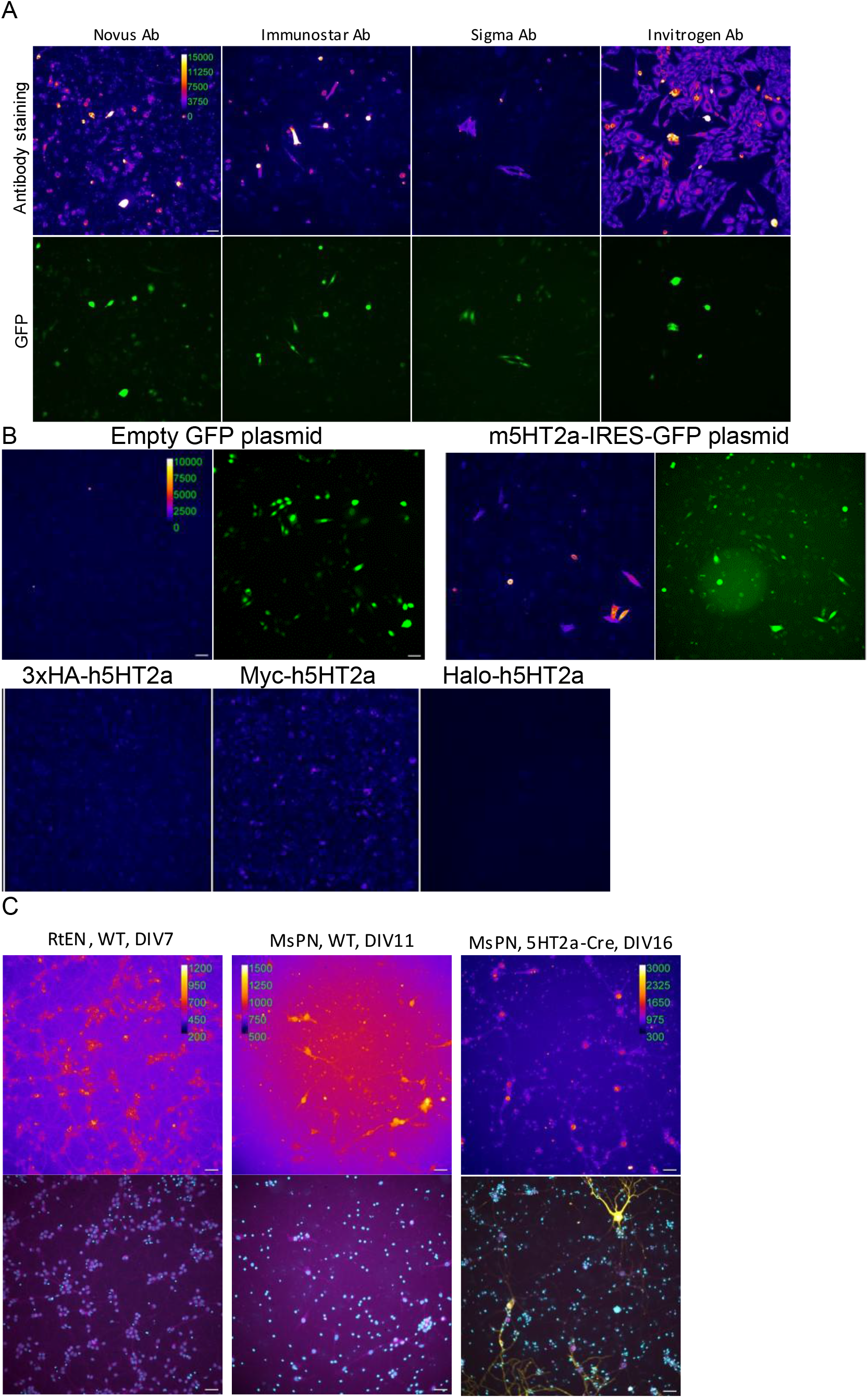
Commercially available antibodies detect over-expressed mouse 5HT2a receptor, but not native receptor in mouse cortical neurons. **A.** 5HT2a receptor antibody screening in CHO-K1 cells transiently transfected with mouse 5HT2a-IRES-GFP. Top row – antibody staining with AF-647, bottom row – GFP expression. Immunostar and Sigma antibodies produced similar membrane staining in GFP positive cells with the visibly least amount of non-specific staining, followed by Novus and Invitrogen Ab. Calibration bar indicating contrast range applicable to all antibody staining images. The scale bar is 40 μm. **B.** Characterization of Immunostar antibody against different 5HT2a receptor constructs and GFP control. Images in “Fire” pseudo color – antibody staining with AF-647, images in green – GFP expression. Immunostar produced 5HT2a specific staining only in CHO cells transiently transfected with mouse 5HT2a receptor, not in cells expressing human 5HT2a receptor. Calibration bar and scale bar is applicable to all images in “Fire” pseudo color. **C.** Immunostar labeling of 5HT2a in RtEN on DIV7, MsPN wild type (WT) on DIV11, and MsPN 5HT2a-Cre on DIV16. Top image – antibody staining with AF-647 for RtEN and MsPN WT and AF-488 for MsPN 5HT2a-Cre in “Fire” pseudo color, bottom image – overlay of nuclear staining (Hoechst, cyan color), antibody staining (magenta), and TdT fluorescence in yellow for MsPN 5HT2a-Cre cultures. Immunostar antibody produced weak non-specific staining, the brightest stained structure being either small puncta around nuclei and in dendrites or cell nuclei.

**Figure S6.**
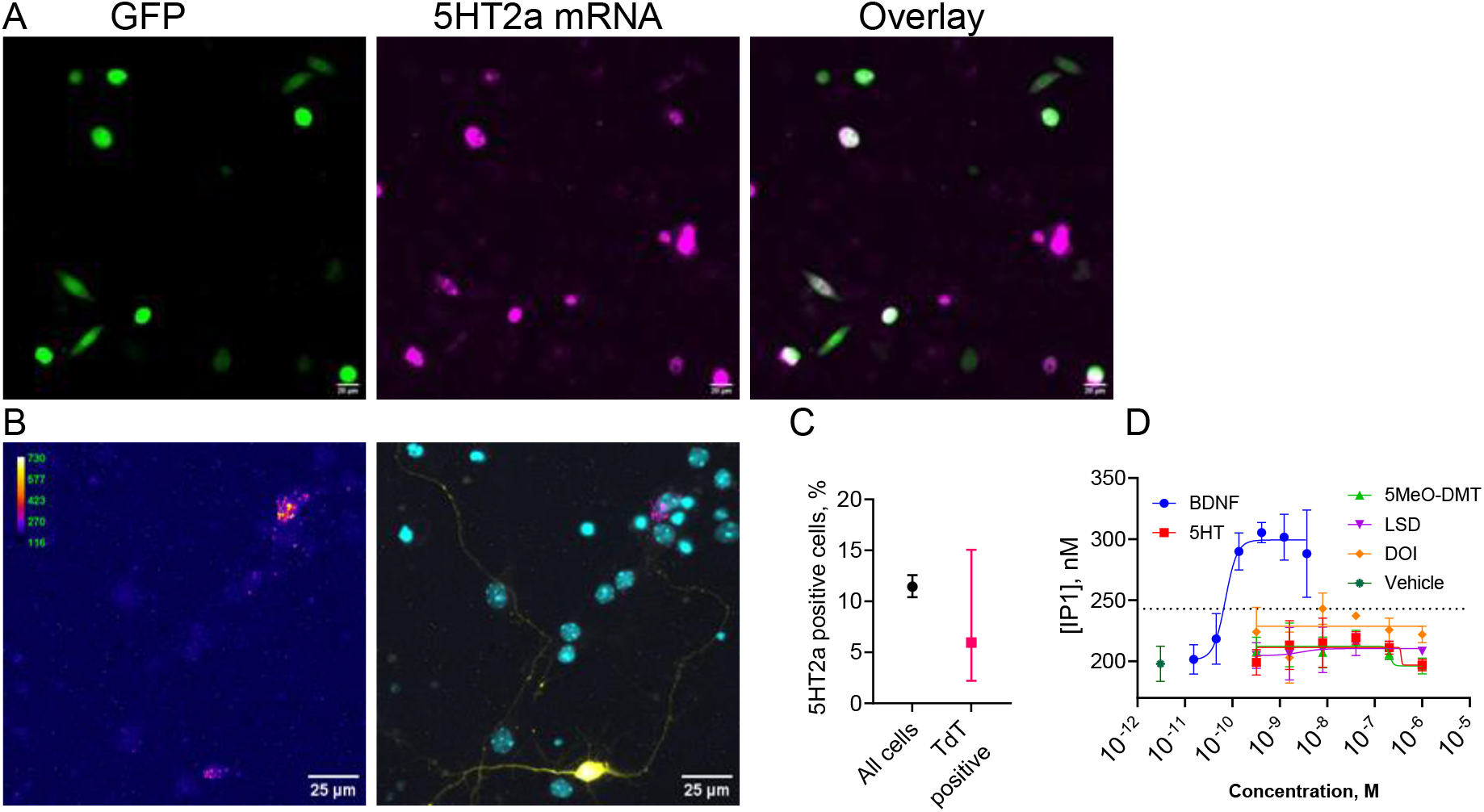
Mouse postnatal (MsPN) cortical neurons at DIV7 have very low levels of 5HT2a receptors in terms of both gene expression and functional receptor protein. **A.** 5HT2a smFISH probe characterization in CHO cells transfected with mouse 5HT2a-IRES-GFP construct. Left image – GFP expression, middle – smFISH probe for mouse 5HT2a mRNA, right – overlay of the images to the left. Scale bar = 20 μm. Probe for mouse 5HT2a mRNA produced bright uniform or puncta staining in GFP positive cells confirming specificity for the 5HT2a mRNA. **B.** Representative images of mouse postnatal cortical cultures treated with vehicle or BDNF. Left image - 5HT2a mRNA smFISH staining, right images – overlayed images of nuclei (cyan, Hoechst staining), a randomly stained neuron (yellow, TdT staining), and 5HT2a mRNA (magenta, same as in left image). **C.** Percent of 5HT2a positive cells in MsPN on DIV7. Cell containing a single smFISH punctum considered to be 5HT2a positive. Data represent mean ± 95% CI of back-transformed estimates derived from the hurdle mixed effect model with negative binomial distribution from 2 μDishes, 10-24 images/plate, 141-371 cells/image. **D.** IP1 accumulation in MsPN induced by BDNF and psychedelics. BDNF significantly increased IP1 concentration in a dose dependent manner (EC50 = 52 nM, or 2 ng/ml), while psychedelics did not. Data represents mean ± SEM of interpolated values for demonstration purpose, nonlinear fit to determine EC50 values were performed on HTRF values.

To measure arborization changes specifically in 5HT2a expressing neurons, we attempted to test MsPN culture derived from 5HT2a-Cre transgenic mice. However, in accordance with smFISH data, only weak TdT expression was observed (data not shown). These results confirmed that like the rat cultures, 5HT2a receptor expression is similarly limited in MsPN cultures, making either system unsuited to assessing cellular responses to psychedelic agents on the timeline compatible with the dendritogenesis assay (DIV3-10).

## Discussion

Interest in psychedelic compounds as treatments for mood disorders, substance use disorders, and other psychiatric and neurological conditions is high and increasing. Currently, the neuroplasticity model – the ability of psychedelic substances to rapidly restore and/or remodel synaptic and circuitry connections and function – is the predominant neurobiological explanation for the rapid and lasting therapeutic effects. Visualization of the morphological changes of cultured rodent cortical neurons, such as dendritogenesis and synaptogenesis, has become a widely used, first-line approach to obtaining the initial evidence of neuroplasticity effects of classic or novel psychedelics. The resulting images of rapidly growing neurons have also acquired a role of simplified visual symbols of complex neuroplasticity phenomena in the broader lay conversations in this area.

Despite the well-known limitations of cultured neurons, they provide useful experimental platforms for detailed probing of molecular signaling mechanisms underlying the morphological and functional changes. While there is strong evidence supporting 5HT2a receptor’s central role in mediating the psychedelic effects in humans and psychedelic-like behaviors in rodents, the molecular link between 5HT2a receptor signaling and neuronal arborization or synaptic morphological remodeling lacks conclusive support^47,53–55^. Assuming the 5HT2a receptor is a relevant primary molecular target in this regard, BDNF release and TrkB activation has been suggested as the downstream mechanism mediating the expression of immediate early genes and other neuroplasticity processes^17,23,56^. A recent alternative hypothesis places 5HT2a receptor outside the neuroplasticity-inducing signaling processes, by claiming that psychedelic compounds, specifically LSD and psilocin, directly interact with TrkB and potentiate its cellular signaling via TrkB dimerization and association with PLCγ, and Erk activation^15^.

Our findings do not support the latter hypothesis as we did not observe any detectable activation of the TrkB signaling cascades induced by the psychedelic compounds either alone (agonistic mode) or in the presence of BDNF (PAM mode). We made the same observations in neuronal cultures endogenously expressing TrkB or CHO cells overexpressing TrkB, as well as in the presence of two of TrkB’s native neurotrophic ligands – BDNF and NT3. These negative results are consistent with our previous attempts to identify and validate small molecule TrkB agonists and PAMs. Despite an extensive effort examining more than 40K drug-like compounds, we did not find any robust, small molecule modulators of TrkB^6^. Collectively, our results do not support the mechanistic model of direct modulation of TrkB by psychedelic compounds.

Regarding the 5HT2a-centered hypothesis, several reports demonstrated that serotonin receptor 2A protein is present in cortical neuronal cultures using immunostaining of the receptor by commercially available antibodies without any quantification of the number of 5HT2a-positive cells^14,17,44,45^. Despite our extensive attempts, we could not confirm detectable protein levels in our cultures due to insufficient specificity/sensitivity of these antibodies. Further, we adapted the IP1 immuno-detection assay in neurons and validated it via BDNF-TrkB-IP3 signaling – only to find no functional expression of the 5HT2a receptor. Instead, we found that the number of cells containing 5HT2a receptor mRNA transcripts was approximately 10% in MsPN cultures.

The previous evidence for psychedelic-induced neuronal arborization in cultured neurons also lacks rigor as the methods are based on snapshot cross-sample comparisons with relatively low sample numbers. We therefore developed an unbiased robust morphological method, MORPHAN, to track dendritic growth over time. Using TdT staining allows us to sparsely label the culture and to accurately trace individual neurons without sacrificing the density and health of the cultures. Moreover, TdT staining allows for visualization of the whole neuron including thin long axons which are often missed using other cellular labeling methods. The within-sample normalization allowed us to develop a powerful statistical analysis to account for random variations of the culture preparation at the single cell level. In the within-sample normalized dataset we never observed significant random effects between the cultures within the same group. On the other hand, MAP2 immuno-staining and between-sample comparison data contained a large random effect of the seeding rendering the treatment effect insignificant despite large sample size. Moreover, pretreatment imaging of the tracked neurons allowed to remove any explicit or implicit biases associated with the selection of the neurons for the analysis and provided greater context for the effects compounds have on neuronal morphology. Additionally, the use of transgenic models would allow conditional expression of TdT and study of dendritogenesis in the specific population of neurons. Coupling it with orthogonal fluorescent labeling could also expand the pool of differentiated neuronal population allowing for multiplex dendritogenesis assay. While we focused on NMax value, there are many more morphological features that could be quantified.

We applied the MORPHAN pipeline in two widely used primary cultures. In RtEN cortical cultures we found equivalent arborization growth in vehicle and BDNF treated samples, masking dendritogenic effects of TrkB signaling, despite its strong functional expression. This result was similar to the mouse neurons cultured in the highQ medium, indicating a strong sensitivity of the cultured neurons to the composition of the medium. Indeed, the standard neurobasal medium was optimized for rat hippocampal cultures to maximize the arborization rate^43^. Hence, to observe dendritogenic effects of BDNF in rat cortical culture it would require preparing the cultures in suboptimal conditions. On the other hand, the standard neurobasal medium appeared to be suboptimal for MsPN cultures, and BDNF treatment rescued this deficiency. In the context of psychedelics and other neuro-therapeutics, however, the expression of key molecules should guide the optimization efforts.

We found no evidence of dendritogenic effects of the psychedelic compounds in mouse cortical cultures. Moreover, 5MeO-DMT and LSD appeared to reduce the growth of neuronal arborization. As discussed above, we found that the number of cells expressing the 5HT2a receptor was limited in MsPN cultures, demonstrating that the standard conditions of primary cultures could not be used to study dendritogenic effects of 5HT2a receptor activation. As such, culturing conditions that drive 5HT2a expression while simultaneously restricting neuronal growth to allow for robust morphological measurements must be developed.

The results we report here provide guidance for future efforts in this active area of research and therapeutics development. For morphological assays to be reliable, attention must be paid to several key experimental factors. Appropriate growth conditions that will neither mask nor exaggerate treatment effects must be determined. The inclusion of positive, negative, and neutral treatment conditions aids in the identification of “basal” conditions for comparison. The method of cell labeling must be carefully considered as not all methods will reveal the same number or the same features of all neurons. A labeling method that provides staining that is sparse enough to allow visual isolation of the neurons but at the same time allows for a cell density that is optimal for the health of the culture and the expression of the protein of interest is particularly important. Methods for imaging, tracing, and measuring neuronal arbor must include measures to prevent biases. Automating these steps not only reduces bias, but it also allows higher throughput and therefore larger sample sizes than can be achieved with manual methods. Within-subject measurements are more robust, as they provide internal normalization, minimizing random effects associated with culturing and handling of the sample, in contrast to snapshot groupwise comparisons. Our careful analytic approach suggests that many previously published studies are critically underpowered, a major issue for reproducibility. Finally, a statistical approach that considers all possible sources of variability and is based on the most appropriate model is essential.

There is growing evidence that psychedelic action engages neuroplasticity to alter brain activity and circuitry. Our data suggests that the manifestation of this neuroplasticity requires specific conditions such as expression of specific molecular targets that is not readily available in widely used young primary cultures. Our results do not dispute that the therapeutic effects of psychedelic treatments are mediated by changes in neuronal plasticity. Instead, our results provide information that will guide researchers in how to develop and use appropriate in vitro models in which to reliably test hypotheses about the role of plasticity in drug therapies, both old and new. We believe thorough characterization of the appropriate neuronal models and development of unbiased, robust, multimodular methods are necessary to uncover the molecular and structural mechanisms of the effects of psychedelics, a class of drugs which offer exciting new avenues for mental health treatments.

## Material and methods

### Materials

BDNF and NT3 were purchased from Peprotech, Serotonin hydrochloride from ThermoFisher, DOI and LSD from Sigma, 5-MeO-DMT from Cayman Chemical, DMSO from Acros Organics. pAAV-FLEX-tdTomato was a gift from Edward Boyden (Addgene viral prep # 28306-PHPeB; http://n2t.net/addgene:28306 ; RRID:Addgene_28306). AAV-CAG-tdTomato (codon diversified) was a gift from Edward Boyden (Addgene viral prep # 59462-PHPeB; http://n2t.net/addgene:59462 ; RRID:Addgene_59462). pENN.AAV.hSyn.Cre.WPRE.hGH was a gift from James M. Wilson (Addgene viral prep # 105553-AAV9; http://n2t.net/addgene:105553 ; RRID:Addgene_105553). 5HT2a-IRES-GFP plasmid was prepared by Vector Biolabs. c-Myc-5-HT2A was a gift from Javier Gonzalez-Maeso (Addgene plasmid # 67944 ; http://n2t.net/addgene:67944 ; RRID:Addgene_67944). 3xHA-5HT2a plasmid was purchased from cDNA Resource Center. pAAV-GFP was a gift from John T Gray (Addgene plasmid # 32395 ; http://n2t.net/addgene:32395 ; RRID:Addgene_32395). List of antibodies is presented in Table S1.

### Animals

All animal studies were approved by the New York State Psychiatric Institute Animal Care and Use Committee and were performed in accordance with PHS Policy on Humane Care and Use of Laboratory Animals. 5HT2a-Cre mice (Tg(Htr2a-cre)KM208Gsat) were a generous gift of Dr. Gergely Turi and bred as Cre x Cre pairs in our animal facility to increase the likelihood of Cre positive pups.

### Primary cultures

E18 rat embryonic cortical tissues were provided by Dr. Clarissa Waites lab or purchased from Transnetyx. Tissues were digested with papain solution prepared in-house using Dr. Sulzer lab protocol or with Miltenyi Neural Tissue Dissociation Kit (130-094-802). Papain solution was prepared by mixing Papain suspension (Worthington Biochemical Corporation) in 1 mM cysteine solution to yield 20 units/ml, then adding H&B 5x concentrate (580 mM NaCl, 27 mM KCl, 130 mM NaHCO3, 10 mM NaH2PO4, 5 mM MgSO4, 2.5 mM EDTA, 125 mM Glucose), sodium kynurenate (final concentration 500 μM), Phenol red (final concentration 0.001%), and 5N HCl to achieve pH7 based on the Phenol Red color. Tissue was placed in papain solution for 15min at 37°C with swirling every 5 min. Digested tissues were sequentially washed twice in Hi En solution (HBSS, 1% BSA, 1% Trypsin Inhibitor, 500 μM sodium kynurenate) and thrice in Lo En solution (HBSS, 0.1% BSA, 0.1% Trypsin Inhibitor, 500 μM sodium kynurenate) to inhibit and remove papain enzyme. Washed and digested tissue were transferred to standard neurobasal medium (NB: neurobasal medium, 2% B27, 0.5 mM Glutamax), triturated, and cell counted. Digestion with Miltenyi Neural Tissue Dissociation Kit was performed according to the manufacturer’s manual. After the digestion, trituration, cells were applied to 70 μm strainer and centrifuged at 150 rcf for 10 min at 4°C and resuspended in NB. The typical yield was 10-12.5 M cells/brain.

P0-P2 mouse postnatal cortical cultures were prepared from 5HT2a-Cre newborn pups (wild type and 5HT2a-Cre cultures) or purchased from Transnetyx (wild type cultures only). For 5HT2a-Cre tissue, newborn pups were chilled on ice for 5 min, decapitated, and heads kept on ice. The brains were extracted on a chilled dish and dissected in HBSS with HEPES (Sigma) on ice. Full cortex was used for seeding. For genotyping, tails of each pup were collected, digested with REDExtract-N-Amp Tissue PCR kit (digestion conditions: 10 min at 55°C followed 3 min 95°C incubation in thermal cycler), and DNA segments amplified through PCR using F-ACTGGGATCTTCGAACTCTTTGGAC and R-GATGTTGGGGCACTGCTCATTCACC primer pair for sample loading control, and F-CCATCTGCCACCAGCCAG and R-TCGCCATCTTCCAGCAGG for Cre transgene using miniAmp thermal cycler (AppliedBiosystems by Thermo Fisher Scientific) and the following protocol: 94°C 3’ → 30x of 94°C 30’’, 60°C 1’, 65°C 1’ + 5’’/cycle → 4°C hold. The amplified DNA was resolved on E-Gel 2% agarose gel (Invitrogen, G401002) on E-Gel Power Snap Electrophoresis device (Invitrogen). The double band between 300-700 bp size indicated Cre-positive tissue. The genotyping was performed during tissue dissection. The Cre-positive tissues were selected for seeding. Tissues were digested for seeding using Miltenyi Neural Tissue Dissociation Kit as described for rat embryonic tissue. To remove the cell debris, cells were washed and centrifuged at 4°C 3-4 times until the supernatant looked transparent. While the precipitate appeared to be drastically reduced in volume, in our experience the lost material contained little amount of intact cell bodies and consisted mostly of cell debris. The typical yield was 2.5-5 M cells/brain. Seeding MsPN cultures without extensive removal of debris resulted in cultures with poor quality and reduced cell survival. Seeding cells in media containing FBS or in Neurobasal medium Plus (Gibco, A3582901) with B27 Plus supplement (Gibco, A3653401) resulted in an increased number of non-neuronal cells. Lipofection of neuronal cultures with Lipofectamine^TM^ 2000 (Invitrogen, 11668019), Lipofectamine^TM^ 3000 (Invitrogen, L3000015), or Effectene® (Qiagen, 301425), were not successful for long term expression of the constructs, as the procedure appeared to be toxic to the cells. AAV infection produced little toxicity but required longer incubation (3-5 days) before the expression of the reporter genes were detectable.

### Cell lines

CHO-K1 cell line was purchased from ATCC (CCL-61) and cultured in Ham’s F-12 medium supplemented with 10% FBS, 2 mM L-Glutamine, and 1% Penicillin-Streptomycin (100 U/ml). For probe validation, CHO cells were transfected with either 5HT2a-IRES-GFP, Myc-5HT2a, 3xHA-5HT2a, or empty-GFP plasmids using Lipofectamine 3000 reagent and a carrier DNA of similar size at 1:3 ratio. The transfection was performed for 24-72h in 6-well plates. CHO-Halo-5HT2a FLP-IN T-Rex cell line was a generous gift of Prof. Jonathan Javitch and Dr. Wesley Asher and cultured in Ham’s F-12 medium supplemented with 10% FBS, 2 mM L-Glutamine, 1% Penicillin-Streptomycin (100 U/ml), 15 μg/ml of Blasticidin, and 500 μg/ml Hygromycin. Receptor expression was induced in the induction medium with 10% dialyzed FBS and 2.22 μg/ml tetracycline for 1-3 days in a 6-well plate. Invitrogen CHO K1 cell line with CellSensor® construct (TrkB-NFAT-bla, K1491) was obtained from Life Technologies and cultured in growth medium (DMEM with GlutaMax) supplemented with 10% dialyzed FBS (Invitrogen, 26400-044), 1x MEM NEAA (Sigma, M7145), 25 mM HEPES (Sigma, H0887), 5 μg/mL Blasticidin (Life Technologies, R21001) and 200 µg/ml Zeocin (Life Technologies, R25001).

### Immunofluorescence

#### 5HT2a Antibody validation

CHO cells expressing 5HT2a receptor (either through transfection or induction with tetracycline) were collected from 6-well plates using MilliporeSigma™ PBS Based Enzyme Free Cell Dissociation Solution (S-014-C), centrifuged at 4°C, 150 rcf, resuspended in growth medium, and seeded at 250k/well on ibidi μ-Slide 8-well coverslip (ibidi, 80806) coated with poly-D-lysine (PDL). RtEN and MsPN cultures were seeded in the same coverslips in standard neurobasal medium. The medium was refreshed on DIV3 and supplemented every 3-5 days to maintain the same medium level. Cell cultures were fixed with 4% formaldehyde (20 min, RT), permeabilized through 3x washing with PBS 0.05% Tween-20 (PBST), and immunostained for 2h at room temperature with anti-5HT2a antibodies from Immunostar (1/200 dilution, rabbit polyclonal, 24288), MilliporeSigma (1/300 dilution, rabbit polyclonal, PC176), Novus Biologicals (1/200 dilution, goat polyclonal, NBP226091), and Invitrogen (1/100 dilution, rabbit polyclonal, PA5-53111) diluted in 3% BSA PBST. After the incubation with primary antibodies, samples were washed 3x 5min with PBST and counter stained for 1h with nuclear stain (Hoechst 33258) and secondary antibodies: rabbit antibodies - goat anti-rabbit conjugated with Alexa Fluor 647 (ThermoFisher, A27040, 1/1000 dilution); goat antibody - rabbit anti-goat conjugated with Alexa Fluor 647 (ThermoFisher, A27018, 1/1000 dilution). After the counter staining, samples were washed 3x 5min with PBST and placed in glycerol mounting medium (20 mM Tris, pH8, 90% glycerol, 0.5% N-propyl gallate) and imaged on epi-fluorescent microscope Leica 4000B equipped with HC PL APO 20x/0.80 objective, PCO Panda camera, Sutter Lambda LS with Xenon lamp.

#### Map2 staining for dendritogenesis assay

RtEN cells were seeded in PDL coated 24-well plates in standard neurobasal medium at 50k/well. The cells were treated with BDNF (50 ng/ml) and Vehicle on DIV5 and fixed on DIV7 using 4% formaldehyde. Immunostaining of MAP2 protein was performed as described above using anti-MAP2 antibody (CellSignaling, #8707, 1/1000 dilution) as primary antibody, anti-rabbit conjugated with Alexa Fluor 488 (CellSingaling, #4412, 1/1000 dilution) as secondary antibody, and Hoechst as a nuclear stain. MsPN cultures used to collect AAV based dataset were fixed on DIV8. Cells were seeded on ibidi μDishes and treated as described in MORPHAN protocol section. The samples were immunostained with the same primary antibody and counterstained with Hoechst and anti-rabbit conjugated with Alexa Fluor 647 as secondary antibody. Samples were imaged with 20x objective in PBST or glycerol mounting medium.

### smFISH

Single molecule fluorescent in-situ hybridization staining was performed using Hybridization Chain Reaction (HCR^tm^) technology from Molecular Instruments^57^. Probes for mouse 5HT2a receptor mRNA were purchased from Molecular Instruments. The staining protocol was based on the manufacturer’s manual for mammalian cells on a slide. All the reagents and equipment were certified RNAse-free. Cell cultures were seeded on ibidi μ-Slide 8-well coverslip or 35 mm μ-Dish with gridded polymer coverslip (81166) coated with PDL. Cells were washed with PBS and fixed with 4% formaldehyde PBS solution for 10-20 min at room temperature. After the fixation, samples were washed twice with ice-cold PBS and permeabilized with 70% ethanol water solution (no buffer) overnight at −20°C. After permeabilization, samples were washed twice with 2X SSC (20x SSC diluted 10 times in mQ water), pre-incubated with manufacturer’s hybridization buffer for 30 min at 37°C, and hybridized with smFISH probes overnight at 37°C. After hybridization, samples were washed 4x for 5 min at 37°C with manufacturer’s wash buffer and twice with 5x SSCT (4 times diluted 20x SSC in mQ water, 0.1% Tween-20) and pre-incubated with manufacturer’s amplification buffer for 30 min at room temperature. In a PCR tube, conjugated to Alexa Fluor 647 h1 and h2 hairpin stock solutions were heated at 95°C in thermal cycler and let cool at RT for 30 min. Hairpin working solution was prepared by diluting hairpin snap-cooled stock solutions in the amplification buffer and applied to the sample. Amplification step was performed overnight at RT in the dark. The excess hairpins were washed off by 2x washing for 5 min at RT with 5xSSCT. Nuclei were stained by 30 min incubation at RT with Hoechst diluted in 5xSSCT. Samples were prepared for imaging by 2x washing for 5 min at RT with 5xSSCT and applying glycerol mounting medium. Imaging was performed on Leica microscope with 20x or 63x oil immersion objective. The number of puncta per cell in neuronal cultures was quantified through CellProfiller^58^ pipeline. Data preparation was performed using Python script and statistical analysis was performed in R. To determine the number of cells expressing 5HT2a receptor and take into account zero-inflated dataset and random effects associated with imaging field-of-view, hurdle mixed effect model with negative binomial distribution from GLMMadaptive library^59^ was applied. Zero-inflated coefficients and upper and lower critical limits estimated through emmeans library were back transformed into percentage through 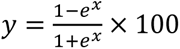 equation, where y is percentage and x - estimated values.

### Molecular signaling assays

#### Culture treatment

For molecular signaling experiments, neuronal cultures and CHO cell lines expressing 5HT2a or TrkB were seeded in PDL coated white clear-bottom 96-well plates. RtEN and MsPN cultures were seeded in standard neurobasal medium at 50k/well in 200 μl/well of medium with no medium exchange until the day of experiments. Induced CHO-Halo-5HT2a FLP-In Trex cells were seeded in induction medium. CHO-TrkB cells were seeded in the respective growth medium. On the day of experiment, medium was exchanged to neurobasal medium (100 μl/well for agonistic mode or 75 μl/well for PAM mode) without any additives in RtEN and MsPN cultures, to Ham’s F12 medium in CHO-Halo-5HT2a cell cultures, and DMEM medium in CHO-TrkB cultures to starve the cells and reduce the background signal. For ELISA and ELFI experiments, cells were starved for 1-3 hours before treatment. For IP1 assay more than 6h incubation was necessary to increase the dynamic range of the readout. Compounds were serially diluted in DMSO (series of 3x, 4x, 5x, or 10x dilutions) and then transferred to experimental medium to yield 200x dilutions in vehicle solution before delivery to the cells (5x dilution, 25 μl/well) to achieve overall 1000x dilution from the stock solution to the cell cultures (0.1% DMSO concentration across the plate). For PAM mode, cells were preincubated for 10 min with the compounds before addition of BDNF or NT3. For IP1 assay, cells were pre-incubated with 3.12 mM LiCl solution for 5 min before addition of compounds to yield 2.5 mM LiCl. For ELISA and ELFI assays, compounds were treated for various time points as indicated in the data. For IP1 assay, cells were treated for 1h before the readout development. Each concentration was tested in triplicates or quadruplicates. The treatment was performed with pseudo randomization with the aim to have each replicate collected from different parts of the plate while minimizing operator’s delivery error.

#### ELISA and ELFI

ELISA to quantify TrkB phosphorylation and ELFI to quantify Erk and Akt phosphorylation were performed as reported previously with no modifications^6^. For ELISA, medium was carefully removed, and cells were lysed on ice with the lysis buffer containing Triton X100, a cocktail of protease (MilliporeSigma, P8340) and phosphatase inhibitors (MilliporeSigma, P5726). The plates were frozen at −80C until the day of ELISA readout. The detection of pTrkB and total TrkB was performed on 2 separate NUNC Immunlon 4 HBX plates as described previously ^6^. For ELFI, cells were fixed with 4% formaldehyde at RT 20 min, permeabilized with PBST, and immunostained for pAkt, Akt, Erk, and pErk through stripping and reprobing protocol described previously ^6^.

#### IP1 assay

IP1 assay kit was adapted for adherent cell cultures. After the treatment, the treatment medium was exchanged without any washing to 1:1 mixture of Neurobasal medium and Stimulation buffer B for RtEN and MsPN cultures, or 1:1 mixture of HAM’s F12 and 1x Stimulation buffer B for CHO-5HT2a, or 1:1 mixture of DMEM and 1x Stimulation buffer B for CHO-TrkB cells (35 μl/well). Immediately, 7.5 μl of the acceptor and donor solutions in the lysis buffer (prepared based on the kit manual) were added sequentially. Cell cultures were incubated at RT for 1h before 2×20 μl of the lysates were transferred to 2 wells in HTRF 384-well low volume opaque white plate (Revvity, 6008280). The time resolved fluorescence of the donor and FRET based emission of the acceptor were recorded using the Synergy Neo2 plate reader (BioTek) using an excitation wavelength of 330/80 nm and emission wavelengths at 620/10 nm and 665/8 nm via Biotek IP1 dual filter cubes (Biotek, 8040505) using the following settings: sensor gain - 129 for ex. / 131 for em., focal height - 4.5 mm, signal collection - 150 μs delay before collecting 50 data points over 500 μsec.

#### Statistical analysis of the molecular signaling datasets

Raw values derived from ELISA and ELFI assays were normalized per well by dividing signal from phosphorylated protein to signal from total protein. For time course experiments, the ratio values were normalized to the Vehicle within the experimental plate. Statistical analysis was performed in GraphPad Prism version 10. The effect of the treatment was tested using two-way ANOVA with Dunnet’s post-hoc multiple comparisons against the vehicle treatment within the time point. To determine EC50 values in dose response dataset, the ratio values were fitted to “[Agonist] vs. response – Variable slope (four parameters)” model with Least square regression method, no constraints or pre-determined initial values or range, with 95% confidence level.

For IP1 assay dataset, the technical replicates were averaged before pooling the data for graph preparation and statistical analysis. To measure EC50 values, the HTRF values were fitted to “[Inhibitor] vs. response – Variable slope (four parameters)” model for IP1 assay with Least square regression method, no constraints or pre-determined initial values of range, with 95% confidence level. Standard curves prepared using the kit manual were used to interpolate HTRF values to IP1 concentration by pooling all the data into one GraphPad data table and fitting the curve with a four-parameter sigmoidal model (Sigmoidal, 4PL). The representative IP1 graphs were created by pooling together each interpolated replicate and fitting “[Agonist] vs. response – Variable slope (four parameters)” model for representation purposes.

### MORPHAN protocol

#### Cell treatment and image acquisition

RtEN and MsPN cells were seeded on ibidi 35 mm μDish with gridded polymer coverslips coated with PDL at 150-300k cells/dish in NB. The cell suspension was added to the insert covered with 500 μl of medium. The cells were allowed to attach overnight in the incubator. On DIV1, AAV PHP.eB particles carrying TdT under CAG promoter at 500 GC/cell multiplicity of infection (MOI) or a pair of AAV9 particles carrying Cre under hSyn promoter construct (MOI 2000) and AAV PHP.eB particles carrying FLEx-TdT under CAG promoter construct (MOI 2000) were added to the cultures, incubated for 2-3h at 37°C before addition of 2 mL of fresh NB. For highQ conditions, MsPN cells were cultured in Neurobasal medium with 2 mM GlutaMax. On DIV3, the medium was changed with 2 mL of fresh neurobasal medium. On DIV5, TdT expressing neurons were imaged using Leica microscope with 20x objective and each plate was given an ID. The same day, another experimenter added compounds to cells. Compounds were prediluted in the fresh neurobasal medium at 50x dilution and then delivered to the designated wells to yield 1000x dilution from the stock. BDNF was added at final concentration of 50 ng/ml, psychedelic compounds, 5HT, DOI, LSD, and 5MeO-DMT, at 1 μM. Solution of neurobasal medium with 0.1% DMSO and 10^−4^% BSA served as vehicle control. The neurons imaged on DIV5 were imaged on DIV6 and DIV7 again by the experimenter blinded to the treatment.

#### Image analysis

##### Background correction

Because of uneven illumination, captured fluorescent images contained a dome-shaped background which hindered the effectiveness of WEKA Machine Learning Segmentation. To mitigate this, we first generated an artificial background using a non-linear regression model in the Python script using the following function:

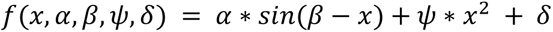

The script read the raw image as an array size of 2048×2048, processing each row to fit the above model to generate a potential background. Since this model estimates were skewed by foreground signal, these regions were excluded through iterative thresholding and a background signal was simulated. Specifically, each image row was iterated upon a number of times defined by a user. During each iteration, the fitted line from the model was normalized, and its mode and interquartile range were used to calculate a threshold that defined the foreground region. Within the region a random value drawn from a Gaussian distribution with the sum of mean and the quarter of the standard deviation of the fitted line as input was assigned to the pixel, generating a simulated background with an artificial noise. The artificial background images were then used in CellProfiler “Correct Illumination - Calculate plugin” with the following settings:

For Correct Illumination – Calculate
Intensity Choice: Regular
Dilate Objects: No
Rescale Option: Yes
(Rescale) Each or All: Each
Smoothing Method: Median
Automatic Object Width: Automatic
Size of Smoothing Filter: 10

Then the function was applied to each image with a “Divide” option in the “Correct Illumination - Apply plugin”.

##### WEKA Machine Learning Segmentation

Background corrected images were pooled into image stacks of 40-60 images and loaded into Trainable WEKA Segmentation plugin v3.3.2^34^ in ImageJ (FIJI)^60^. Segmentation settings were as follows: training features – gaussian blur, difference of gaussians, mean, maximum, median, Kuwahara filter; minimum sigma – 1, maximum sigma – 16; classifier – random forest, batch size 100, number of features 2, number of trees 200; 3 classes for background, soma, and projections. Blinded operators trained model until the classifier produced satisfactory segmentation at the discretion of the operators. Image outliers that were poorly segmented by the model were segmented using a new model. Each stack was segmented using newly trained models.

##### Manual correction and matching

We found that most of the neurons would unpredictably migrate on the surface within the imaging time frame. Thus, we could not develop an automated approach to match the same neurons imaged on different days. To this end, we created a Python script allowing for visual matching of the neurons by the experimenters blinded to the treatment. We also recognized that WEKA segmentation contained disconnected area within the perceived projections because of the low signal in the original images. To correct this, the experimenter manually checked and corrected each image to improve the accuracy of the automated tracing using ImageJ.

##### APP2 Vaa3D

Automatic tracings of neurons were generated using the APP2 algorithm from Vaa3D^36,61^ in a Python script with the WEKA segmented binary images. First, an operator identified somas positions using the ImageJ tools. We then developed a Python script to process the saved data by matching the ROI coordinates to their respective images. Second, we developed another Python script to run the APP2 algorithm for each image prepared in a headless mode. The algorithm was sped up with a multiprocessing package in the script. The APP2 algorithm settings were as follows:

Background Threshold: Auto
Auto Downsample: 0
Radius from 2D: 0
Gray Scale Distance Transform: 0
Allow Gap: 0
Length Threshold: 5
Signal Redundancy Threshold: 0.5
CNN type: 1
High Intensity: 0

The output of the Python script is a tracing file with .swc extension.

##### G-Cut

For images with a cluster of neurons that grew on top of each other, the APP2 algorithm considered the touching neurons as one single object, producing one single tracing file. To address this issue, we used G-Cut^37^ to separate and re-organize the tracings. The G-Cut algorithm required a tracing file in a .swc format and a set of points in pixels that represented the somas in the cluster. Since APP2 plugin produced tracings with slightly different coordinates for soma positions than were identified prior, the mismatch caused an error in G-Cut. To avoid this error, we created a Python script that automatically searched for the nearest coordinates in the tracing file from the originally identified soma positions. The preprocessed set of data was then used as input for the G-Cut algorithm with the following settings: z_scale=5; conditional=mouse; and cell_type=neocortex. For each APP2 tracing of any given image, all soma coordinates within the image were provided.

##### Dendrogram and rectified Sholl plot generation

The single neuron tracing files were first preprocessed to extract the lengths, the branching points, and parent-child relationships of neurites in ImageJ Macro. The extracted data was then used to reconstruct the neuron with straightened neurites. The reconstructed images were structured from left to right and top to bottom such that axons are placed on the far left of images and the child branches are placed thereafter. For better visualization, a child branch with the longest distance from the root of the parent branch was placed immediately to the right of that parent branch. Once a neuron was reconstructed, a rectified Sholl plot was generated by counting the number of branches per row of the reconstructed image from top to bottom.

##### NMax measurement

Data for statistical analysis was prepared using Python notebooks. As a measurement of neuronal arborization, the maximal number of crossings (NMax) on the rectified Sholl plot within 150 μm from soma was detected using a custom Python function. The plot was smoothed using a median filter of 10 um and peaks were found within the distance limit indicated in the function from SciPy library^62^.

#### Statistical analysis

The NMax values were not normally distributed and represented a count dataset violating assumptions of t-test and ANOVA. Moreover, t-test and ANOVA could not account for intra- and inter-experimental variations. We examined different models to find an appropriate statistical test^42,63^. We examined standard unpaired t-test and nested t-test of raw and log-transformed data in GraphPad, linear model of raw and log-transformed data, generalized linear with negative binomial and Poisson distributions, linear mixed effect model of raw and log-transformed data with each dish or well as random effect, and generalized mixed effect model with negative binomial and Poisson distributions and each dish or well as random effect in R using lme4 library^64^. For the unpaired data we found the generalized mixed effect model with negative binomial to produce the most aligned Q-Q plot and normal distribution of residuals. For the paired data, we found generalized mixed effect model with Poisson distribution to be the most appropriate model.

Data analysis was performed using R. The derived NMax values were fitted to generalized mixed effect model with negative binomial distribution, NMax as function of treatment, each dish or well as random for unpaired data and Poisson distribution, NMax as function of interaction of time and treatment, each neuron as random effect for paired data of lme4 package. The difference in the interaction of time and treatment was used to test the significant effect of the treatment. To determine the growth difference within the treatment for paired data, the model was post-hoc analyzed using emmeans package^65^ with defined comparison pairs and Sidak correction for multiple comparisons. The confidence intervals on the graphs were estimated using bootstrapping method of the model estimates and back transformed through y = e^x^ function, where x – estimate, y – back transformed value.

### Data handling and code accessibility

Collected images were stored on OMERO server^66^. The data analysis was performed using Jupyterlab notebooks. The analysis pipeline for MORPHAN will be publicly available at https://github.com/Hyun-P/MorphAn.

## Supplementary Materials

### 5HT2a receptors antibody validation

We screened four available antibodies, including Immunostar antibody specificity of which was confirmed in 5HT2a KO mice^2^. To examine the specificity of the antibodies we initially used CHO-K1 cells transiently transfected with mouse 5HT2a-IRES-GFP construct in which GFP expression should correspond to 5HT2a expression. Immunostar antibody produced visible contrast between GFP positive and negative cells with little non-specific staining. Interestingly, Immunostar and Sigma antibodies produce identical staining patterns. Closer examination of antibody sources indicated that these antibodies could be identical. Immunostar antibody did not stain human 5HT2a, however. We reasoned that the antigen region of human and mouse proteins were different enough to preclude antibody binding to the human 5HT2a. Applying Immunostar to MsPN cortical cultures on DIV11 did not produce the expected membrane or cytosolic staining. Instead, there seemed to be nuclear non-specific staining with autofluorescence of the sample. This result indicated that expression of 5HT2a may not be prominent in these cultures.

**Table S1.** List of antibodies used in this study.

| **Target** | **Vendor** | **Cat. #** | **Host** |
| --- | --- | --- | --- |
| **Pan-pTyr** | R&D Systems | HAM1676 | Mouse IgG<sub>1</sub> |
| **TrkB** | R&D Systems | AF397 | Goat pAb |
|  | Sino Biologicals | 10047-RP02 | Rabbit pAb |
| **5HT2a** | Immunostar | 24288 | Rabbit pAb |
|  | MilliporeSigma | PC176 | Rabbit pAb |
|  | Novus | NBP226091 | Goat pAb |
|  | Invitrogen | PA5-53111 | Rabbit pAb |
| **p-AKT (pS473)** | Cell Signaling | #4060 | Rabbit mAb |
| **AKT1** | Cell Signaling | #2938 | Rabbit mAb |
| **p-ERK** | Cell Signaling | #4370 | Rabbit mAb |
| **ERK** | Cell Signaling | #9102 | Rabbit pAb |
| **MAP2** | Cell Signaling | #8707 | Rabbit Ab |
| **Rabbit IgG** | Cell Signaling | #7074 | Goat Ab |
| **Rabbit IgG** | Cell Signaling | #4412 | Goat Ab |
| **Rabbit IgG** | ThermoFisher | A27040 | Goat Ab |
| **Goat** | ThermoFisher | A27018 | Rabbit Ab |

## Acknowledgements

We would like to thank Dr. Clarissa Waites, Columbia University for providing us with rat embryonic cortical tissues, Dr. Jonathan Javitch and Dr. Wesley Asher, Columbia University/New York State Psychiatric Institute for providing us with 5HT2a receptor expressing cell lines and plasmids. We would like to thank Dr. David Sulzer, Dr. Eugene Mosharov, and Dr. Se Jun Choi, Columbia University/New York State Psychiatric Institute for providing instruction and insight on neuronal culturing and experimental conditions. We would like to thank Diane Lu, Dr. Leon Fernandes, and Claire Ming-Yi He, Columbia University for assistance with the statistical analysis. We would also like to thank Firuz Mukhitdinov for technical support of the OMERO server and remote workstation. We would also like to thank Dr. Frances Zakusilo and George Zakusilo for invaluable feedback on project development. U.B. and D.S. discloses support for the research of this work from Mather Foundation [grant number MF-2001-00633].

## Author contributions

U.B., D.S., and E.H.S. conceptualized the project and manuscript. U.B. and H.P. designed the experiments and MORPHAN pipeline. U.B., H.P., K.B., M.D., K.H, E.A. performed MORPHAN. H.P., U.B., M.B., and J.W. developed codes. U.B. performed statistical analysis. U.B., K.B., J.W., B.D., M.B., K.H., and N.G. performed ELFI, ELISA, and IP1 assays. U.B., M.D., E.A., J.W., M.B., and N.G. performed immunostaining and smFISH. The manuscript was drafted by U.B., D.S., and E.H.S., and edited by all authors. All authors agreed with the manuscript in its current form.

## References

1. Pittenger, C. & Duman, R. S. Stress, depression, and neuroplasticity: a convergence of mechanisms. Neuropsychopharmacology 33, 88–109 (2008).

2. Krystal, J. H. et al. Ketamine and the neurobiology of depression: Toward next-generation rapid-acting antidepressant treatments. Proc Natl Acad Sci U S A 120, e2305772120 (2023).

3. Licznerski, P. & Duman, R. S. Remodeling of axo-spinous synapses in the pathophysiology and treatment of depression. Neuroscience 251, 33–50 (2013).

4. Levy, M. J. F. et al. Neurotrophic factors and neuroplasticity pathways in the pathophysiology and treatment of depression. Psychopharmacology vol. 235 2195–2220 Preprint at 10.1007/s00213-018-4950-4 (2018).

5. Kim, J., He, M. J., Widmann, A. K. & Lee, F. S. The role of neurotrophic factors in novel, rapid psychiatric treatments. Neuropsychopharmacology 2023 49:1 49, 227–245 (2023).

6. Boltaev, U. et al. Multiplex quantitative assays indicate a need for reevaluating reported small-molecule TrkB agonists. Sci Signal 10, (2017).

7. de Vos, C. M. H., Mason, N. L. & Kuypers, K. P. C. Psychedelics and Neuroplasticity: A Systematic Review Unraveling the Biological Underpinnings of Psychedelics. Front Psychiatry 12, 724606 (2021).

8. Béïque, J. C., Imad, M., Mladenovic, L., Gingrich, J. A. & Andrade, R. Mechanism of the 5-hydroxytryptamine 2A receptor-mediated facilitation of synaptic activity in prefrontal cortex. Proc Natl Acad Sci U S A 104, 9870–9875 (2007).

9. Vollenweider, F. X. & Kometer, M. The neurobiology of psychedelic drugs: implications for the treatment of mood disorders. Nature Reviews Neuroscience 2010 11:9 11, 642–651 (2010).

10. Bender, D. & Hellerstein, D. J. Assessing the risk–benefit profile of classical psychedelics: a clinical review of second-wave psychedelic research. Psychopharmacology 2022 239:6 239, 1907–1932 (2022).

11. Goodwin, G. M. et al. Single-Dose Psilocybin for a Treatment-Resistant Episode of Major Depression. New England Journal of Medicine 387, 1637–1648 (2022).

12. Carhart-Harris, R. et al. Trial of Psilocybin versus Escitalopram for Depression. New England Journal of Medicine 384, 1402–1411 (2021).

13. Kadriu, B. et al. Ketamine and Serotonergic Psychedelics: Common Mechanisms Underlying the Effects of Rapid-Acting Antidepressants. International Journal of Neuropsychopharmacology 24, 8–21 (2021).

14. Vargas, M. V et al. Psychedelics promote neuroplasticity through the activation of intracellular 5-HT2A receptors. Science 379, 700–706 (2023).

15. Moliner, R. et al. Psychedelics promote plasticity by directly binding to BDNF receptor TrkB. Nat Neurosci 26, 1032–1041 (2023).

16. Ilchibaeva, T. et al. Serotonin Receptor 5-HT2A Regulates TrkB Receptor Function in Heteroreceptor Complexes. Cells 2022, Vol. 11, Page 2384 11, 2384 (2022).

17. Ly, C. et al. Psychedelics Promote Structural and Functional Neural Plasticity. CellReports 23, 3170–3182 (2018).

18. Reichardt, L. F. Neurotrophin-regulated signalling pathways. Philos Trans R Soc Lond B Biol Sci 361, 1545–64 (2006).

19. Masson, J., Emerit, M. B., Hamon, M. & Darmon, M. Serotonergic signaling: Multiple effectors and pleiotropic effects. Wiley Interdiscip Rev Membr Transp Signal 1, 685–713 (2012).

20. Kim, K. et al. Structure of a Hallucinogen-Activated Gq-Coupled 5-HT2A Serotonin Receptor. Cell 182, 1574–1588.e19 (2020).

21. Watts, S. W. Activation of the Mitogen-activated Protein Kinase Pathway via the 5-HT2A Receptora. Ann N Y Acad Sci 861, 162–168 (1998).

22. Jaggar, M. & Vaidya, V. A. 5-HT2A receptors and BDNF regulation: Implications for psychopathology. Receptors 32, 395–438 (2018).

23. Vaidya, V. A., Marek, G. J., Aghajanian, G. K. & Duman, R. S. 5-HT 2A Receptor-Mediated Regulation of Brain-Derived Neurotrophic Factor MRNA in the Hippocampus and the Neocortex. The Journal of neuroscience : the official journal of the Society for Neuroscience vol. 17 http://www.ncbi.nlm.nih.gov/pubmed/9092600 (1997).

24. Lee, F. S. & Chao, M. V. Activation of Trk neurotrophin receptors in the absence of neurotrophins. Proc Natl Acad Sci U S A 98, 3555–60 (2001).

25. Iwakura, Y., Nawa, H., Sora, I. & Chao, M. V. Dopamine D1 receptor-induced signaling through TrkB receptors in striatal neurons. Journal of Biological Chemistry 283, 15799–15806 (2008).

26. Mitre, M. et al. Transactivation of TrkB Receptors by Oxytocin and Its G Protein-Coupled Receptor. Front Mol Neurosci 15, 891537 (2022).

27. Ly, C. et al. Transient Stimulation with Psychoplastogens Is Sufficient to Initiate Neuronal Growth. ACS Pharmacol Transl Sci 4, 452–460 (2021).

28. Dunlap, L. E. et al. Identification of Psychoplastogenic N, N-Dimethylaminoisotryptamine (isoDMT) Analogues through Structure-Activity Relationship Studies. J Med Chem 63, 1142–1155 (2020).

29. Lewis, V. et al. A non-hallucinogenic LSD analog with therapeutic potential for mood disorders. Cell Rep 42, (2023).

30. Langhammer, C. G. et al. Automated Sholl analysis of digitized neuronal morphology at multiple scales: Whole cell Sholl analysis versus Sholl analysis of arbor subregions. Cytometry Part A 77 A, 1160–1168 (2010).

31. Ji, Y. et al. Acute and gradual increases in BDNF concentration elicit distinct signaling and functions in neurons. Nature Neuroscience 2010 13:3 13, 302–309 (2010).

32. Studer, L. et al. Effects of brain-derived neurotrophic factor on neuronal structure of dopaminergic neurons in dissociated cultures of human fetal mesencephalon. Exp Brain Res 108, 328–336 (1996).

33. Kellner, Y. et al. The BDNF effects on dendritic spines of mature hippocampal neurons depend on neuronal activity. Front Synaptic Neurosci 6, 5 (2014).

34. Arganda-Carreras, I. et al. Trainable Weka Segmentation: a machine learning tool for microscopy pixel classification. Bioinformatics 33, 2424–2426 (2017).

35. Peng, H., Ruan, Z., Long, F., Simpson, J. H. & Myers, E. W. V3D enables real-time 3D visualization and quantitative analysis of large-scale biological image data sets. Nat Biotechnol 28, 348–353 (2010).

36. Xiao, H. & Peng, H. APP2: automatic tracing of 3D neuron morphology based on hierarchical pruning of a gray-weighted image distance-tree. Bioinformatics 29, 1448–1454 (2013).

37. Li, R. et al. Precise segmentation of densely interweaving neuron clusters using G-Cut. Nature Communications 2019 10:1 10, 1–12 (2019).

38. Aliaga Maraver, J. J., Mata, S., Benavides-Piccione, R., DeFelipe, J. & Pastor, L. A method for the symbolic representation of neurons. Front Neuroanat 12, 1–19 (2018).

39. Schierwagen, A. Neuronal morphology: Shape characteristics and models. In Neurophysiology vol. 40 310–315 (Springer, 2008).

40. Bird, A. D. & Cuntz, H. Dissecting Sholl Analysis into Its Functional Components. Cell Rep 27, 3081–3096.e5 (2019).

41. Arshadi, C., Günther, U., Eddison, M., Harrington, K. & Ferreira, T. SNT: A Unifying Toolbox for Quantification of Neuronal Anatomy. bioRxiv 2020.07.13.179325 (2020) doi:10.1101/2020.07.13.179325.

42. Wilson, M. D., Sethi, S., Lein, P. J. & Keil, K. P. Valid statistical approaches for analyzing sholl data: Mixed effects versus simple linear models. J Neurosci Methods 279, 33–43 (2017).

43. Brewer, G. J., Torricelli, J. R., Evege, E. K. & Price, P. J. Optimized survival of hippocampal neurons in B27-supplemented neurobasal^TM^, a new serum-free medium combination. J Neurosci Res 35, 567–576 (1993).

44. Desouza, L. A. et al. The Hallucinogenic Serotonin2A Receptor Agonist, 2,5-Dimethoxy-4-Iodoamphetamine, Promotes cAMP Response Element Binding Protein-Dependent Gene Expression of Specific Plasticity-Associated Genes in the Rodent Neocortex. Front Mol Neurosci 14, 1–15 (2021).

45. Xia, Z., Hufeisen, S. J., Gray, J. A. & Roth, B. L. The PDZ-binding domain is essential for the dendritic targeting of 5-HT2A serotonin receptors in cortical pyramidal neurons in vitro. Neuroscience 122, 907–920 (2003).

46. González-Maeso, J. et al. Identification of a serotonin/glutamate receptor complex implicated in psychosis. Nature 452, 93–97 (2008).

47. González-Maeso, J. et al. Hallucinogens Recruit Specific Cortical 5-HT2AReceptor-Mediated Signaling Pathways to Affect Behavior. Neuron 53, 439–452 (2007).

48. Salahpour, A. et al. BRET biosensors to study GPCR biology, pharmacology, and signal transduction. Front Endocrinol (Lausanne) 3, 30991 (2012).

49. Olsen, R. H. J. et al. TRUPATH, an open-source biosensor platform for interrogating the GPCR transducerome. Nature Chemical Biology 2020 16:8 16, 841–849 (2020).

50. Pottie, E. & Stove, C. P. In vitro assays for the functional characterization of (psychedelic) substances at the serotonin receptor 5-HT2AR. J Neurochem 162, 39–59 (2022).

51. Trinquet, E. et al. d-myo-Inositol 1-phosphate as a surrogate of d-myo-inositol 1,4,5-tris phosphate to monitor G protein-coupled receptor activation. Anal Biochem 358, 126–135 (2006).

52. Gong, S. et al. A gene expression atlas of the central nervous system based on bacterial artificial chromosomes. Nature 425, 917–925 (2003).

53. Kwan, A. C., Olson, D. E., Preller, K. H. & Roth, B. L. The neural basis of psychedelic action. Nature Neuroscience 2022 25:11 25, 1407–1419 (2022).

54. Shao, L.-X. et al. Psilocybin induces rapid and persistent growth of dendritic spines in frontal cortex in vivo. Neuron 109, 2021.02.17.431629 (2021).

55. Hesselgrave, N., Troppoli, T. A., Wulff, A. B., Cole, A. B. & Thompson, S. M. Harnessing psilocybin: Antidepressant-like behavioral and synaptic actions of psilocybin are independent of 5-HT2R activation in mice. Proc Natl Acad Sci U S A 118, 1–7 (2021).

56. González-Maeso, J. et al. Transcriptome fingerprints distinguish hallucinogenic and nonhallucinogenic 5-hydroxytryptamine 2A receptor agonist effects in mouse somatosensory cortex. J Neurosci 23, 8836–43 (2003).

57. Choi, H. M. T., Beck, V. A. & Pierce, N. A. Next-generation in situ hybridization chain reaction: Higher gain, lower cost, greater durability. ACS Nano 8, 4284–4294 (2014).

58. Stirling, D. R. et al. CellProfiler 4: improvements in speed, utility and usability. BMC Bioinformatics 22, 1–11 (2021).

59. Rizopoulos D. GLMMadaptive: Generalized Linear Mixed Models using Adaptive Gaussian Quadrature. (2023).

60. Schindelin, J. et al. Fiji: an open-source platform for biological-image analysis. Nature Methods 2012 9:7 9, 676–682 (2012).

61. Peng, H., Bria, A., Zhou, Z., Iannello, G. & Long, F. Extensible visualization and analysis for multidimensional images using Vaa3D. Nat Protoc 9, 193–208 (2014).

62. Virtanen, P. et al. SciPy 1.0: fundamental algorithms for scientific computing in Python. Nature Methods 2020 17:3 17, 261–272 (2020).

63. Warton, D. I., Lyons, M., Stoklosa, J. & Ives, A. R. Three points to consider when choosing a LM or GLM test for count data. Methods Ecol Evol 7, 882–890 (2016).

64. Bates, D., Mächler, M., Bolker, B. M. & Walker, S. C. Fitting Linear Mixed-Effects Models Using lme4. J Stat Softw 67, 1–48 (2015).

65. Searle, S. R., Speed, F. M. & Milliken, G. A. Population marginal means in the linear model: An alternative to least squares means. American Statistician 34, 216–221 (1980).

66. Allan, C. et al. OMERO: flexible, model-driven data management for experimental biology. Nature Methods 2012 9:3 9, 245–253 (2012).

